# TCP3-mediated regulation of cell expansion in *Arabidopsis thaliana*

**DOI:** 10.1101/2025.08.24.672037

**Authors:** Tomotsugu Koyama, Tadashi Kunieda, Hiromi Toyonaga, Mika Nobuhara, Nobutaka Mitsuda, Kouichi Soga, Junko Ishida, Motoaki Seki, Koji Takahashi, Toshinori Kinoshita, Ayumu Bessho, Taku Demura, Masaru Ohme-Takagi

## Abstract

(1) Cell expansion is crucial for organ morphogenesis in multicellular organisms. Apoplast acidification triggers plant cell expansion. Plant hormones and transcription factors such as TEOSINTE BRANCHED, CYCLOIDEA, and PROLIFERATING CELL NUCLEAR ANTIGEN BINDING FACTORs (TCPs) control cell expansion. However, details regarding the regulatory mechanism of cell expansion for organ morphogenesis remain unclear.
(2) In this study, we used molecular, biochemical, cellular, genetic, atomic force microscopy, and tensile testing analyses and showed that miR319-targeted TCPs integrated cell expansion with organ morphogenesis in *Arabidopsis thaliana*.
(3) We found that TCPs directly induce the expression of genes encoding cell wall loosening proteins and SMALL AUXIN UP RNAs (SAURs), activators of plasma membrane–localized H^+^-ATPases. TCP-mediated activation of plasma membrane– localized H^+^-ATPases stimulates apoplast acidification, reduce cell stiffness, promote cell expansion, and thus exaggerate elongation of the hypocotyl. Ectopic expression of a *SAUR* gene in sextuple *tcp* mutant plants substantially recovered the hypocotyl morphology of the *tcp* mutant, providing genetic evidence of *TCP*-mediated *SAUR* regulation.
(4) Collectively, our data show that TCPs regulate apoplast acidification for cell expansion.

## Introduction

Cell expansion is essential for plant organ morphogenesis and growth. The coordinated regulation of cell division and expansion underlies the morphogenesis of leaves, whereas the regulation of cell expansion determines the extent of hypocotyl elongation (Hepworth & Lenhard, 2014; Zuch *et al*., 2022; Krahmer & Fankhauser, 2024; Schneider *et al*., 2024). Under stress conditions such as high salt concentrations, cell expansion is limited to enable plants to cope with the stress, as turgor-driven growth consumes water (Geilfus, 2017; Rui & Dinneny, 2020). Plant cells are surrounded by a rigid cell wall, which is a barrier for cell expansion (Chebli & Geitmann, 2017; Cosgrove, 2023). The acid growth theory hypothesizes that apoplast acidification allows cells to expand due to turgor pressure (Rayle & Cleland, 1992; Kutscheras, 1994; Arsuffi & Braybrook, 2018; Du *et al*., 2020).

Plasma membrane (PM)-localized H^+^-ATPases are responsible for apoplast acidification and cell expansion (Haruta *et al*., 2015). The activation of PM H^+^-ATPases requires phosphorylation of the conserved penultimate threonine residue (Hayashi *et al*., 2010; Takahashi *et al*., 2012). The activation of PM H^+^-ATPases by auxin and brassinosteroid (BR) and its repression by abscisic acid (ABA) illustrate the close association between plant hormones and cell expansion (Miao *et al*., 2022). At the PM, auxin activates TRANSMEMBRANE KINASE1, which directly activates PM H^+^-ATPases and thereby enhances hypocotyl cell expansion (Li *et al*., 2021; Lin *et al*., 2021; Friml *et al*., 2022). In the nucleus, auxin induces the expression of the *SMALL AUXIN UP RNA19 (SAUR19)* and *SAUR63* genes, which activate PM H^+^-ATPases by inhibiting protein phosphatases (Spartz *et al*., 2014; Nagpal *et al*., 2022). BR also induces *SAUR* genes and activates PM H^+^-ATPases (Minami *et al*., 2019). Conversely, ABA represses the activity of PM H^+^-ATPases and suppresses hypocotyl elongation (Hayashi *et al*., 2014). In contrast to the plant hormone–mediated regulation of PM H^+^-ATPases, details regarding the mechanisms that connect PM H^+^-ATPases with upstream signaling regulators of organ morphogenesis remain elusive.

Several classes of transcription factors (TFs) link cell expansion with organ morphogenesis (Sarvepalli *et al*., 2019; Krahmer & Fankhauser, 2024). It has been reported that TEOSINTE BRANCHED, CYCLOIDEA, PROLIFERATING CELL NUCLEAR ANTIGEN BINDING FACTOR (TCP) TFs play a key role in the regulation of cell expansion and organ morphogenesis (Nicolas & Cubas, 2016; Lan & Qin, 2020; Rath *et al*., 2022). Among 24 *TCP* genes of *Arabidopsis thaliana* (Arabidopsis), the phylogenetically related *CINCINNATA*-like *TCPs* (designated here as *TCPs* for simplicity) consist of five miR319-targeted genes: *TCP2*, *TCP3*, *TCP4*, *TCP10*, and *TCP24*, as well as non-targeted *TCP5*, *TCP13*, and *TCP17* (Nath *et al*., 2003; Koyama *et al*., 2007). Suppression of *TCP* genes inhibits cell expansion in leaves and cotyledons, accompanied by the formation of wavy and serrated leaves as well as epinastic cotyledons (Paul Alvarez *et al*.; Palatnik *et al*., 2003; Koyama *et al*., 2007, 2010; Efroni *et al*., 2008; Challa *et al*., 2021). Even though *TCP* genes are thought to link auxin metabolism with cell expansion (Challa *et al*., 2016), the precise pathway by which TCPs activate H⁺-ATPase activity and induce apoplast acidification remains insufficiently understood.

While cell division and expansion are distinct processes, it is technically challenging to precisely track the transition from proliferative to differentiative growth during leaf development. In contrast, during cotyledon and hypocotyl morphogenesis, growth becomes primarily driven by cell expansion following an initial phase of cell division. (Cendreau *et al*.; Tsukaya *et al*., 1994; Boron & Vissenberg, 2014). Overexpression of *TCP* genes enhances hypocotyl elongation and suppresses cotyledon epinasty (Palatnik *et al*., 2003; Koyama *et al*., 2007; Challa *et al*., 2016). Thus, cotyledons and hypocotyls could be suitable organs for examining the mechanism of cell expansion governed by *TCP* genes. In this study, we first demonstrated that *TCP3*, a model *TCP* gene for genetic studies, promotes cell expansion in cotyledons and hypocotyls. We then determined that TCP3 decreases cell stiffness. We further showed that TCP3 directly regulates genes encoding SAURs and cell wall loosening proteins, activates PM H^+^-ATPases, and induces apoplast acidification. Finally, we obtained genetic evidence demonstrating *TCP*-mediated *SAUR* regulation. Our study thus provides mechanistic insights into TCP3-regulated cell expansion for organ morphogenesis.

## Materials and methods

### Plant materials and growth conditions

*Arabidopsis thaliana* Col-0 was used throughout the study. Details regarding *Pro35S:mTCP3*, *tcp_s* (*tcp3/4/5/10/13/17*), and *ProSAUR65:GUS* plants were described previously (Koyama *et al*., 2007, 2010, 2017). To generate *ProSAUR20:GUS tcp_s*, *ProSAUR20:GUS* was sequentially crossed with *tcp3, tcp3/4*, *tcp3/4/10*, *tcp3/4/10/13*, and *tcp_s* (Koyama *et al*., 2007, 2010, 2017). Arabidopsis seeds were sterilized using sodium hypochlorite solution, stored at 4°C for 3 days, and then sown on a plate containing half-strength MS salts under a cycle of 16-h light (photosynthetic photon flux density of 50 μmol m^−2^ s^−1^)/8-h dark at 22°C unless otherwise indicated. For application of estradiol to plants in liquid culture, 10-day-old seedlings were transferred into liquid medium containing half-strength MS salts, 5 g/L sucrose, and 0.5 g/L MES and incubated for 36 h with gentle agitation under continuous light conditions, after which compounds that included DMSO and 5 µM estradiol (Wako, Japan) were added to the liquid medium.

### Construction of plasmids

To generate the *ProXVE:mTCP3* plasmid, the coding sequence (CDS) of *mTCP3* was inserted into *pER8* (Zuo et al., 2000; Koyama et al., 2007). To generate the *gTCP3* plasmid, the 3178-bp sequence upstream and 2726-bp sequence downstream were inserted into *pCR8* (Thermo Fisher Scientific, MA). To generate *gTCP3:GUS*, *gTCP3:TCP3-HA*, and *gTCP3:GFP-SAUR20*, the CDSs of *GUS*, *TCP3-HA*, and *GFP-SAUR20* were individually inserted into *gTCP3-pCR8*. To generate *ProEXPA5:GUS*, the upstream sequence of *EXPA5* was inserted into *pGUS_ENTR*. *gTCP3:GUS*, *gTCP3:TCP3-HA*, *gTCP3:GFP-SAUR20*, and *ProEXPA5:GUS* were individually transferred into the plant transformation vector *pBCKH* or *pCBKK* (Mitsuda *et al*., 2006). To generate *ProSAUR20:GUS* and *ProBGAL2:GUS*, the upstream sequence was inserted into *pBI101* (Clontech). Mutation of the TCP-binding sites of the promoters was carried out using PCR with appropriate primer sets (Supporting Information Table **S1**). For EMSAs, the CDS for the N-terminal 108 amino acid residues of TCP3 was amplified using appropriate primers and cloned into *pGEX-6P* (Cytiva, Japan).

### Measurement of cell size

The cotyledons and hypocotyls of 7-day-old seedlings were cleaned with ethanol and transparent solution (choral hydrate 8 g, water 2 mL, and glycerol 1 mL). Hypocotyl cells and cotyledon adaxial pavement cells were observed using differential interference contrast microscopy (Axio Imager A2, Zeiss, Germany). To measure cell size, images of the hypocotyl and pavement cells were analyzed using ZEN2 Pro software (Zeiss). The border lines between cells in Figure **1e** were detected and processed using ImageJ software (Schindelin *et al*., 2012).

**Figure 1.**
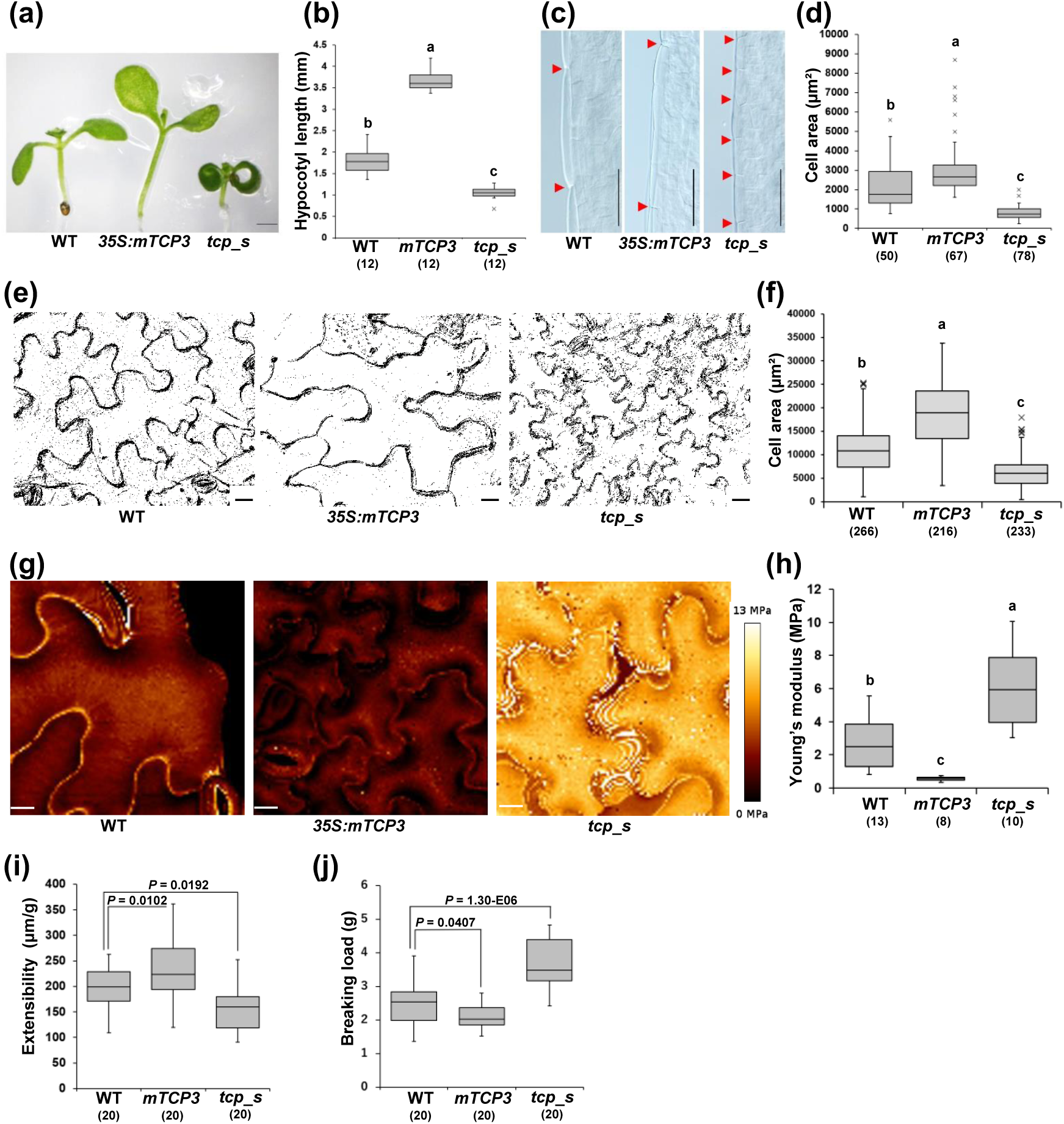
Cell expansion and stiffness regulated by TCP3. (a) Images of 7-day-old seedlings of WT, *Pro35S:mTCP3*, and *tcp_s.* Bar = 1 mm. (b) Hypocotyl length of WT, *Pro35S:mTCP3*, and *tcp_s*. Different letters above box plots indicate statistically significant differences determined using the Tukey test (n = 12). (c) Hypocotyl cells of WT, *Pro35S:mTCP3*, and *tcp_s*. Bars = 100 µm. (d) Size of hypocotyl cells of WT, *Pro35S:mTCP3*, and *tcp_s*. Sample numbers are indicated in parentheses. Different letters above the box plots indicate statistically significant differences determined using the Tukey-Kramer test. (e) Adaxial pavement cells of WT, *Pro35S:mTCP3*, and *tcp_s* cotyledons. Bars = 20 µm. (f) Size of pavement cells of WT, *Pro35S:mTCP3*, and *tcp_s* cotyledons. Sample numbers are indicated in parentheses. Different letters above the box plots indicate statistically significant differences determined using the Tukey-Kramer test. (g) AFM images of the adaxial pavement cells of WT, *Pro35S:mTCP3*, and *tcp_s* cotyledons. The color code of Young’s modulus is presented to the right of the images. Scale bars = 10 µm. (h) Quantification of the Young’s modulus of WT, *Pro35S:mTCP3*, and *tcp_s* cells. Sample numbers are indicated in parentheses. Different letters above the box plots indicate statistically significant differences determined using the Tukey-Kramer test. (i) Cell wall extensibility of WT, *Pro35S:mTCP3*, and *tcp_s* hypocotyls. (j) Breaking load of WT, *Pro35S:mTCP3*, and *tcp_s* hypocotyls. Sample numbers are indicated in parentheses and *P* values were calculated using Dunnett’s test in **(i)** and **(j)**.

### Atomic force microscopy (AFM) analysis

Cotyledons of 7-day-old seedlings were placed on a plastic petri dish and fixed with biocompatible glue (JPK Bio-Compatible Glue, Bruker). For AFM analysis, the adaxial side of the cotyledons was scanned in water at room temperature (turgid conditions) using a NanoWizard 4 microscope (JPK Instruments, Germany) equipped with a CellHesion module (JPK Instruments) and a biosphere B500-NCH cantilever equipped with a spherical tip (Nanotools, Germany). AFM analyses were conducted in the Quantitative Imaging mode with a Setpoint force of 500 nN to acquire force curves over a 100-µm square area with a resolution of 100 × 100 pixels with a speed of 75 µms^−1^ using JPK SPMControl software. The average indentation depth was approximately 40 nm.

To examine cell stiffness, Young’s modulus was calculated according to a Hertz model using data processing software (JPK Instruments). Young’s modulus values were averaged among five regions of interest per pavement cell, excluding the edge region of the cells based on the height images. A total of 8 to 13 cells from 4 to 6 different cotyledons per genotype were used for comparative analysis of cell stiffness.

### Measurement of cell wall extensibility and breaking load

Seedlings were grown for 7 days on agar plates with half-strength MS salts under a 16 h light / 8 h dark cycle at 15 μmol m⁻² s⁻¹. This low-light condition promoted hypocotyl elongation and provided samples of adequate length for tensile testing. Cell wall extensibility and breaking load were measured using a tensile tester (Tensilon STB-1225S; A&D, Tokyo, Japan) following the method described by Hattori *et al*. (2022). The measurement was conducted on a 0.5-mm region of the hypocotyl, located approximately 0.5 mm below the shoot apical meristem. Cell wall extensibility was calculated when the applied load increased from 0.8 g to 1.0 g.

### Gene expression analysis

Total RNA was isolated from frozen plant tissues using Trizol (Thermo Fisher Scientific) or Sepazol (Nacalai Tesque, Japan), cleared by precipitation using 4 M LiCl solution (final concentration), and treated with DNase I (Qiagen, Germany). For RT-PCR analyses, aliquots of total RNA were subjected to reverse transcription using SuperScript III (Thermo Fisher Scientific) with an Oligo(dT) primer (Thermo Fisher Scientific) and analyzed using a CFX96 real-time PCR system (Bio-Rad, CA) using appropriate sets of primers (Supporting Information Table **S1**). A standard curve based upon a reference sample was used to confirm the correct amplification efficiency of the primer pairs and calculate the transcript levels of genes of interest. Relative transcript levels were normalized to that of *UBQ1*. Gene expression analyses were conducted using six biological replicates.

Total RNA was extracted from *ProXVE:mTCP3* plants after application of DMSO or estradiol for 16 h, for which Cy3- and Cy5-labeled cDNAs were prepared, respectively (n = 3), followed by hybridization using an Agilent Arabidopsis 2 Oligo Microarray kit (Agilent Technologies Inc.). All microarray experiments and initial data output were performed according to the supplier’s manual using the feature extraction and image analysis software (version A.6.1.1; Agilent Technologies Inc.). A flag filter provided with the feature extraction software was used to remove data meeting the following criteria: the spot was signal saturated, the signal was non-homogenously distributed, the signal was not significantly greater than background, or the background-subtracted signal was less than the background signal plus 2.6-fold of the standard deviation of the background signal. All signal values were divided by the median of all filter-passed values in each channel to calculate fold-change in gene expression. *P* values were calculated using a *t*-test for paired data. To estimate the false-discovery rate, *Q* values (Storey 2002) were calculated using R statistical software with the qvalue package. We observed that *Q* values were <0.019 when *P* values were <0.05; therefore, selection of differentially expressed genes based on *P*<0.05 was considered to pose no risk of including false-positive identifications. The raw and processed data were deposited in NCBI GEO under accession number GSE271710.

### Chromatin immunoprecipitation (ChIP) assay

Nuclear extracts were prepared from 3-week-old Arabidopsis plants, as described previously (Koyama *et al*., 2010, 2013). The extracts were sonicated and then immunoprecipitated using an anti-HA antibody (clone 3F10; Roche). The chromatin precipitate was reverse cross-linked, purified with ethanol, and used as a template for PCR analysis (CFX96 real-time PCR system; Bio-Rad) using an appropriate set of primers (Supporting Information Table **S1**). Values were calculated based on a standard curve generated from the input sample; values for samples processed without the antibody were set at 1.

### Electro-mobility shift assays (EMSA)

The N-terminal 108 amino acid sequence of TCP3 fused with glutathione S-transferase (GST) was expressed in *Escherichia coli* and purified using a GST SpinTrap (Cytiva) according to the manufacturer’s instructions. To generate probes, Cy5-labeled forward oligo DNAs were annealed with corresponding reverse oligo DNAs. A total of 1 µL of each probe (25 µM) was incubated with GST (3 µg) or GST-TCP3 at room temperature for 30 min, electrophoresed on a 7.5% acrylamide gel, and detected using an Amersham Imager 600 (Cytiva). Non-labeled WT and mutant nucleotide competitors were added to the reaction mixtures. The oligo DNAs used in EMSAs are listed in Supporting Information Table **S1**.

### Detection of GUS activity

Seven-day-old seedlings were fixed in 90% acetone for 10 min on ice, washed with water, and incubated at 37°C for overnight in GUS staining buffer containing 20 mM sodium phosphate (pH 7.0), 20 mg/mL 5-bromo-5-chloro-3-indolyl-β-glucuronide (Nacalai Tesque), 3 mM K_4_[Fe(CN)_6_], 3 mM K_3_[Fe(CN)_6_], and 1% (v/v) DMSO. The samples were then cleared with ethanol and transparent solution described above and observed using an Axio Imager A2 microscope (Zeiss).

### Determination of H^+^-ATPase phosphorylation levels

Seven-day-old seedlings were kept in the dark for 24 h before harvesting to suppress light-induced activation of H^+^-ATPase (Okumura *et al*., 2016). To monitor H^+^-ATPase phosphorylation, protein extracts of dark-adapted plants were prepared as described previously (Takahashi *et al*., 2012), separated using SDS-PAGE on 5-20% (w/v) gradient acrylamide gels, transferred onto nitrocellulose membranes (Amersham Protran, 0.45 µm, Sytiva), and subjected to immunoblotting analysis as described previously (Hayashi *et al*., 2010). The following antibodies were used: anti–H^+^-ATPase (1:3000) (Hayashi *et al*., 2010), anti–phosphorylated Thr_947_ of AHA2 (1:3000) (Hayashi *et al*., 2010), and anti– rabbit IgG, HRP-Linked F (ab’) 2 fragment, donkey (1:7500, Sytiva). Immunoreactive proteins were detected using an ECL prime Western blotting kit (Sytiva) and quantified using an Amersham Imager 600. Levels of H^+^-ATPase phosphorylation were calculated based on the ratio of signal intensity of phosphorylated Thr_947_ of AHA2 to the signal intensity of H^+^-ATPase (Takahashi *et al*., 2012).

### LiCl and fusicoccin (FC) assays

For LiCl assays, 14-day-old seedlings were transferred onto half-strength MS salt plates with or without 20 mM LiCl supplementation and incubated for the indicated number of days. Chlorophyll content was measured using CCM-300 (Opti-Sciences, Hudson).

For FC assays, 3-day-old plants were transferred onto half-strength MS salt plates supplemented with either 1 µM FC or ethanol and grown for 4 days. Images of plants were acquired using a stereomicroscope (SMZ 745T, Nikon).

### 8-Hydroxypyrene-1,3,6-trisulfonic acid trisodium salt (HPTS) staining

HPTS staining was performed as described by Barbez et al. (Barbez et al., 2017) with some modifications. In brief, 4-day-old seedlings were incubated for 1 h in the dark in half-strength MS salt solution (pH 5.7) supplemented with 1 mM HPTS and 0.01% Triton X-100 and then mounted on a slide with a cover glass. Images of fluorescent signals from the protonated HPTS form (excitation, 405 nm; emission peak, 514 nm) and deprotonated HPTS form (excitation, 488 nm; emission peak, 514 nm) were obtained using a FV3000 confocal microscope (Olympus, Japan) with a 20× air objective. Ratiometric images were calculated using a previously described Fiji macro (Barbez et al., 2017). For pH calibration, WT seedlings were incubated in buffer solutions adjusted to pH 3.7–6.7 (20 mM MES, supplemented with 1/2 MS salts and 0.01% Triton X-100), and fluorescence ratios (488/405) were recorded under each condition. Calibration curves were generated by fitting the data using Excel. To avoid extrapolation beyond the calibrated range, pH values below 3.7 and above 6.7 were excluded. Using the derived calibration equations, fluorescence ratios were converted into absolute pH values. Means of ratiometric values were obtained from cells of four cotyledons and six hypocotyls per genotype, and the experiments were repeated three times with similar results.

### Statistical analysis

Statistical analysis and box-plot development were performed using BellCurve_for_Excel software (https://bellcurve.jp/ex/) with the default settings. A Student’s t, Kolmogorov-Smirnov, Tukey-Kramer, or Bonferroni test was performed as indicated in each figure legend. *P* < 0.05 was considered statistically significant.

## Results

### Cell expansion and stiffness regulated by TCP3

To overcome the miRNA-mediated regulation of *TCP* genes and their partial functional redundancy (Supplemental information Figure **S1**), we previously generated transgenic *Arabidopsis* plants that overexpress mutated *TCP3* in the miR319-target sequence (*Pro35S:mTCP3*) for gain-of-function analysis and sextuple *tcp* mutants (*tcp_s*) with genetic mutations in *TCP3*, *TCP4*, *TCP5*, *TCP10*, *TCP13*, and *TCP17* for loss-of-function analysis (Koyama *et al*., 2007, 2017). In this study, to clarify details of cell expansion regulated by *TCP* genes, we analyzed the hypocotyls and cotyledons of *Pro35S:mTCP3*, *tcp_s*, and wild-type (WT) plants. Light microscopic observations showed that, compared with the WT, *Pro35S:mTCP3* enhanced hypocotyl elongation, whereas *tcp_s* suppressed elongation (Figure **1a** and **1b**) (Koyama *et al*., 2007, 2017). To investigate whether the changes in WT, *Pro35S:mTCP3*, and *tcp_s* hypocotyl growth were associated with differential cell expansion, we measured the volume of the hypocotyl cells. Quantitative analysis indicated that, compared with WT hypocotyls, *Pro35S:mTCP3* hypocotyls exhibited larger cell size and *tcp_s* hypocotyls reduced cell size (Figure **1c** and **1d**). Similarly, compared with WT cotyledons, the cell size of *Pro35S:mTCP3* cotyledons was increased, whereas that of *tcp_s* cotyledons was reduced (Figure **1e** and **1f**). These results demonstrate that TCP3 promotes cell expansion.

Enhanced cell expansion is often associated with decreasing cell stiffness (Peaucelle *et al*., 2015; Wu *et al*., 2020). To examine the mechanical properties of cells, AFM analyses have been developed (Milani *et al*., 2011; Peaucelle *et al*., 2011). In our study, the AFM analyses revealed that in comparison with WT cells, *Pro35S:mTCP3* cells had a decreased elastic modulus, whereas the elastic modulus of *tcp_s* cells was increased (Figure **1g** and **1h**). To complement these results, we conducted tensile testing using hypocotyls (Hattori *et al*. 2022). Compared with WT, *Pro35S:mTCP3* hypocotyls exhibited increased cell wall extensibility, whereas *tcp_s* hypocotyls displayed decreased extensibility (Figure **1i**). In contrast, compared with WT, *Pro35S:mTCP3* hypocotyls showed a reduced breaking load, while *tcp_s* hypocotyls exhibited an increased breaking load (Figure **1j**). These tensile data are consistent with the AFM results and suggest that TCP3 may affect both the stiffness and extensibility of cell wall.

### TCP3 downstream genes related to cell wall loosening proteins and apoplast acidification

To investigate how TCP3 regulates genes involved in cell expansion, we performed a transcriptome analysis to identify its downstream targets. The exaggerated hypocotyl elongation phenotype of *Pro35S:mTCP3* might have indirect effects on the transcriptome, and the low fertility of this mutant could make it difficult to conduct reproducible experiments (Koyama *et al*., 2007, 2017). To avoid such problems, we applied a LexA-VP16-estrogen receptor (XVE) system (Zuo et al., 2000) to generate estradiol-inducible *mTCP3* (*ProXVE:mTCP3*) plants. *ProXVE:mTCP3* plants exhibited enhanced hypocotyl elongation in the presence of estradiol for 7 d (Figure **2a**), demonstrating that the estradiol-inducible *mTCP3* gene is functional in planta. To identify the genes downstream of *TCP3*, we performed transcriptome analysis of *ProXVE:mTCP3* plants following application of DMSO or estradiol for 16 h, the duration of which induced expression of the *mTCP3* gene (Supporting Information Figure **S2**). The transcriptome analysis identified 1552 upregulated genes (fold-change ≥1.5 and *P*≤0.05 [*Q* value (Storey, 2002) ≤0.019]) after the application of estradiol (Figure **2b**). Among these genes, we found 181 genes that overlapped with the genes downregulated in *ProXVE:TCP3-SRDX*, in which the dominant negative *TCP3-SRDX* gene was induced by estradiol (Koyama *et al*., 2010), and these genes were consequently categorized as TCP3 downstream genes (Figure **2b**).

**Figure 2.**
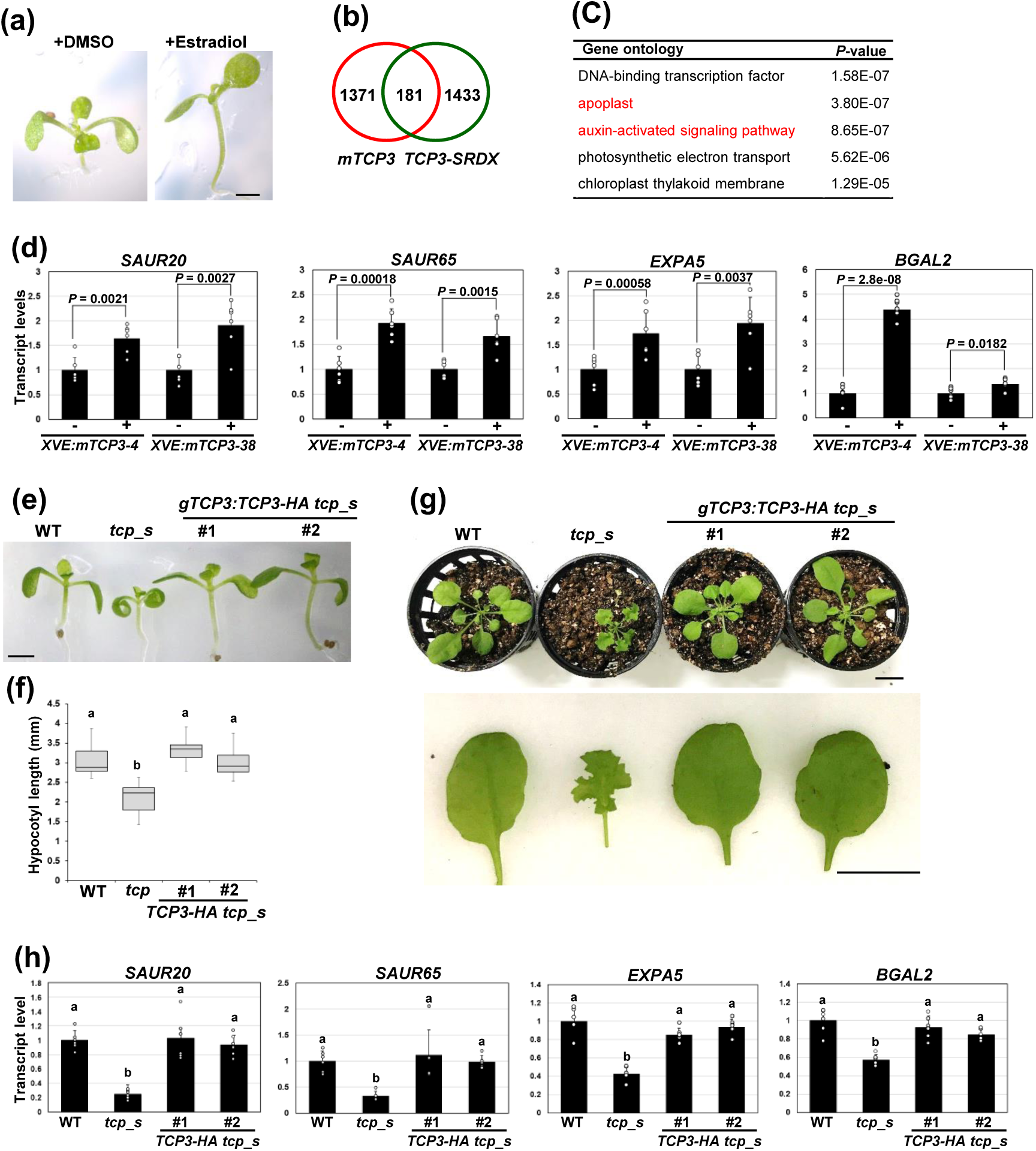
Analysis of the expression of *TCP3* downstream genes. (a) Phenotype of *ProXVE:mTCP3* grown with DMSO or estradiol for 7 days. Bar = 1 mm. (b) Number of genes induced by *mTCP3* (red) and *TCP3SRDX* (green) (Koyama *et al*., 2010). (c) Functional categories of *TCP3* downstream genes enriched according to GO analysis. (d) Relative transcript levels of *TCP3* downstream genes. mRNA samples were prepared from plants supplemented with DMSO (−) or estradiol (+) for 16 h. Two independent *ProXVE:mTCP3* lines were analyzed. Error bars indicate standard deviations of six biological replicates; *P* values were calculated using Student’s *t*-test. (e) Seedling morphology of WT, *tcp_s*, and two independent lines of *gTCP3:TCP3-HA tcp_s*. Bar = 1 mm. (f) Hypocotyl length of WT, *tcp_s*, and two independent *gTCP3:TCP3-HA tcp_s* lines. Different letters above box plots indicate statistically significant differences determined using the Tukey test (n = 18). (g) Rosette (upper panel) and leaf (lower panel) morphology of WT, *tcp_s*, and two independent *gTCP3:TCP3-HA tcp_s* lines. Bar = 1 cm. (h) Relative transcript levels of *TCP3* downstream genes in WT, *tcp_s*, and two independent *gTCP3:TCP3-HA tcp_s* lines. Error bars indicate standard deviations of six biological replicates, and different letters above the bars indicate statistically significant differences determined using the Tukey test.

To explore the biological processes and regulatory pathways affected by TCP3, we performed Gene Ontology (GO) enrichment analysis of the TCP3 downstream genes. GO analysis ranked DNA-binding TFs at the top of the enriched gene group list (Figure **2c**, Supporting Information Figure **S3**). The TF group included *LONG HYPOCOTYL IN FAR-RED (HFR*) (Fairchild *et al*., 2000), *SHORT HYPOCOTYL2/INDOLE-3-ACATIC ACID3 (SHY2/IAA3*) (Tian and Reed 1999), *PHYTOCHROME-INTERACTING FACTOR3 (PIF3)* (Ni *et al*., 1998), *GIBBERELLIC ACID INSENSITIVE (GAI)* (Cowling and Harberd, 1999), *REPRESSOR OF ga1-3 LIKE1 (RGL1)* (Li *et al*., 2016), and *BR-ENHANCED EXPRESSION3 (BEE3)* (Cifuentes-Esquivel *et al*., 2013), which regulate hypocotyl elongation (Supporting Information Figure **S3**). These results are consistent with the phenotype of TCP3-mediated hypocotyl elongation (Figure **1a** and **1b**).

We next focused on gene groups related to apoplast functions and auxin signaling, both of which are closely linked to cell expansion. Importantly, the GO analysis ranked the apoplast and auxin-regulated signaling pathways as second and third among the enriched gene groups, respectively (Figure **2c**). The apoplast group genes encode apoplast-localized cell wall loosening proteins such as alpha-xylosidase 1 (Shigeyama *et al*., 2016), beta-galactosidase 2 (BGAL2) (Gantulga *et al*., 2008), xyloglucan endotransglucosylase/hydrolase 6 (Rose *et al*., 2002), expansin A5 (EXPA5), and EXPA6 (Goh *et al*., 2012) (Figure **2c** and Supporting Information Figure **S4a**). The auxin-activated signaling pathway group genes include INDOLE-3-ACETIC ACIDs (IAAs), AUXIN RESPONSE FACTORs (ARFs), and SAURs (Figure **2c** and Supporting Information Figure **S4b**). Enrichment analysis of the TCP3 downstream genes ranked the SAUR domains at the top among conserved protein domains (Supporting Information Figure **S5a**). *mTCP3* and *TCP3-SRDX* regulated the expression of most *SAUR19* and *SAUR63* subfamily members (Supporting Information Figure **S5b**) (Spartz *et al*., 2014; Nagpal *et al*., 2022). These results demonstrate the close association of TCP3 with cell wall loosening proteins and SAURs. Because cell wall loosening proteins are active under acidic apoplast conditions and SAURs activate PM H^+^-ATPases responsible for apoplast acidification (Spartz *et al*., 2014; Arsuffi & Braybrook, 2018; Du *et al*., 2020), we hypothesized that TCP3 promotes the transcription of genes for cell wall loosening proteins and apoplast acidification and thus regulates cell expansion.

We then analyzed the expression of TCP3 downstream genes. Because we previously proposed *SAUR20* and *SAUR65* as TCP3 downstream genes (Koyama *et al*., 2010), we selected *SAUR20* and *SAUR65* as representatives of the *SAUR19* and *SAUR63* subfamilies, respectively (Spartz *et al*., 2014; Nagpal *et al*., 2022). We also selected *EXPA5* and *BGAL2* as representatives of genes encoding cell wall loosening proteins because they include putative TCP-binding sequences in their promoters (see below). RT-PCR analysis demonstrated the upregulation of *SAUR20* (line 4; 1.64-fold, line 38; 1.91-fold), *SAUR65* (line 4; 1.92-fold, line 38; 1.66-fold), *EXPA5* (line4; 1.73-fold, line 38; 1.93-fold), and *BGAL2* (line 4; 4.39-fold, line 38; 1.37-fold) following the application of estradiol in two independent transgenic *ProXVE:mTCP3* lines (Figure **2d**).

We further investigated whether the transcript levels of *SAUR20*, *SAUR65*, *EXPA5*, and *BGAL2* are decreased in *tcp_s* and recover to normal levels following the introduction of *TCP3* into *tcp_s*. To this end, we introduced *TCP3-HA* under control of the upstream and downstream genomic regions of *TCP3* (*gTCP3*) into *tcp_s* (*gTCP3:TCP3-HA tcp_s,* Supporting Information Figure **S6a** and **S6b**). Compared with the short hypocotyl and highly wavy leaves of *tcp_s*, two independent lines of *gTCP3:TCP3-HA tcp_s* had normal hypocotyls and younger leaves similar to those of WT plants (Figure **2e** to **2g**, see also Supporting Information Figure **S7** for older leaf morphology). RT-PCR analysis indicated that *tcp_s* decreased the expression of *SAUR20* (0.25-fold), *SAUR65* (0.33-fold), *EXPA5* (0.43-fold), and *BGAL2* (0.57-fold), whereas two independent *gTCP3:TCP3-HA tcp_s* lines retained these gene transcripts at similar levels to those in WT plants (Figure **2h**). These results demonstrate that *TCP3* regulates the expression of the *SAUR20*, *SAUR65*, *EXPA5*, and *BGAL2* genes.

### Direct regulation of SAUR20, SAUR65, EXPA5, and BGAL2 by TCP3

We also examined whether TCP3 directly targets *SAUR20*, *SAUR65*, *EXPA5*, and *BGAL2*. The *SAUR20*, *SAUR65*, *EXPA5*, and *BGAL2* promoters harbor putative TCP-binding sequences (Figure **3a**). ChIP assays demonstrated enrichment of the *SAUR20*, *SAUR65*, *EXPA5*, and *BGAL2* promoter sequences using an anti-HA antibody immunoreactive to TCP3-HA in *gTCP3:TCP3-HA tcp_s* (Figure 3A). The negative control *UBQ10* sequence was not enriched in the ChIP assay (Supporting Information Figure **S7**). EMSAs detected the shifted bands associated with complexes of the *SAUR20*, *SAUR65*, *EXPA5*, and *BGAL2* promoters with GST-fused TCP3, but not with GST alone (Figure **3b**). WT nucleotide competitors reduced the intensity of the shifted bands, but the mutated competitors retained the bands (Figure **3b**), indicating specific binding of the *SAUR20*, *SAUR65*, *EXPA5*, and *BGAL2* promoters to TCP3. Promoter-GUS analysis presented strong signals for *ProSAUR20:GUS*, *ProSAUR65:GUS*, *ProEXPA5:GUS*, and *ProBGAL2:GUS* in cotyledons and hypocotyls in which *gTCP3:GUS* was active (Figure **3c**, Supporting Information Figure **S9**). In contrast, the mutated promoters in the TCP-binding sequences, namely, *ProSAUR20m:GUS*, *ProSAUR65m:GUS*, *ProEXPA5m:GUS*, and *ProBGAL2m:GUS*, did not present signals in cotyledons and hypocotyls (Figure **3d**). Additionally, *ProSAUR20:GUS* was crossed to *tcp_s*, and GUS signals in seedlings and leaves of the resultant *ProSAUR20:GUS tcp_s* declined (Figure **3e** and **3f**). These results indicate that TCP3 directly regulates the *SAUR20*, *SAUR65*, *EXPA5*, and *BGAL2* genes.

**Figure 3.**
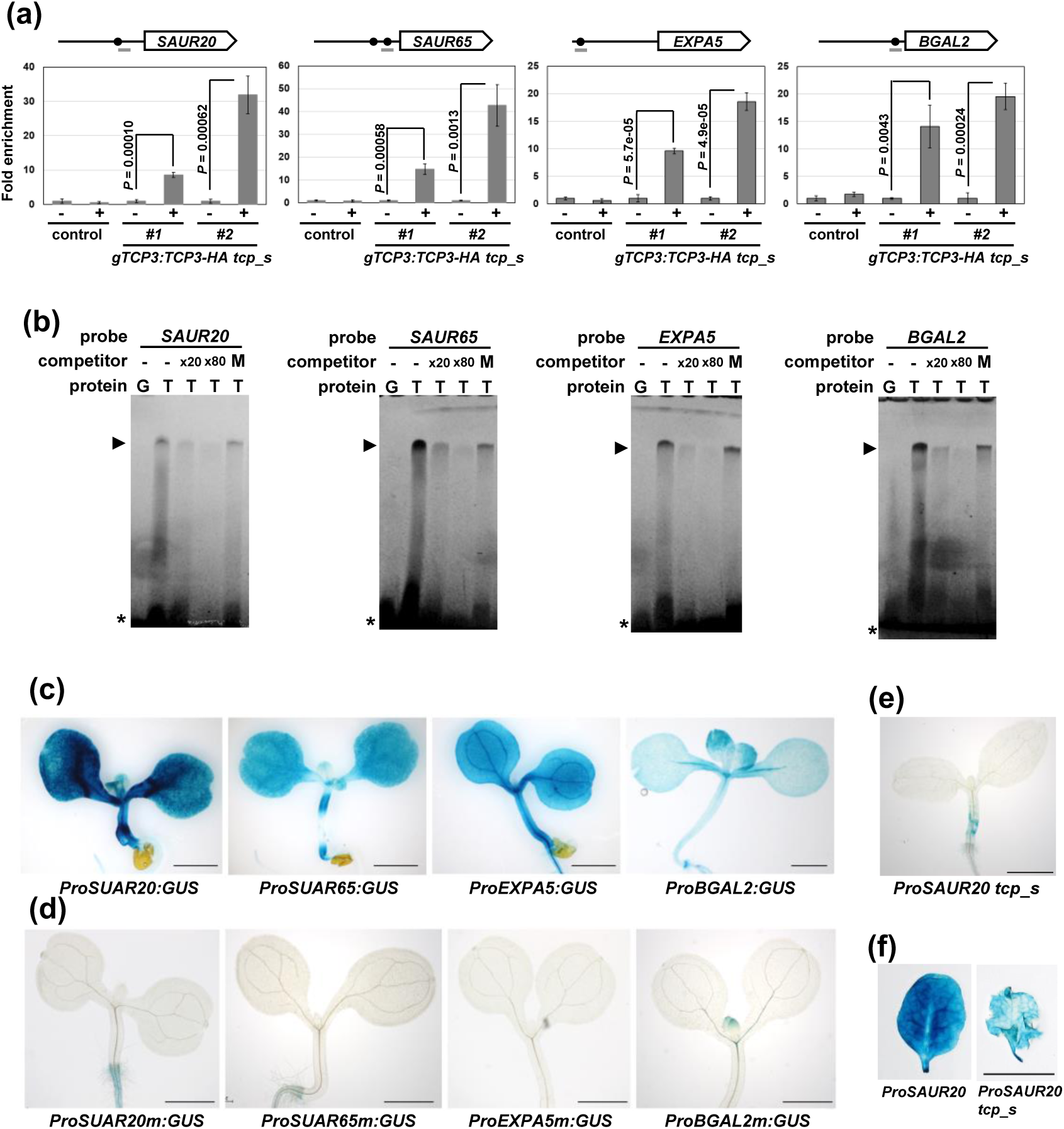
Direct regulation of TCP3 downstream genes. (a) ChIP analysis of TCP3 downstream genes. The diagrams above the graphs present the relative regions of amplicons (gray bars) around the TCP-binding site (black circle) that are located at −354 bp of *SAUR20*, −126 and −258 bp of *SAUR65*, −2334 bp of *EXPA5*, and −76 bp of *BGAL2*. The chromatin of *tcp_s* (control) and two independent *gTCP3:TCP3-HA tcp_s* lines was immunoprecipitated with (+) or without (−) an anti-HA antibody, and the precipitated DNA fragments were quantified using real-time PCR. Fold-enrichment was determined relative to the value determined in the absence of antibody. Error bars indicate the standard deviation of technical triplicates. Graphs show a representative result of three biological replicates with similar trends. (b) EMSAs showing specific binding of the promoter sequences with TCP3. Labeled probes were incubated with GST (G) or GST-TCP3 (T) in the absence (−) or presence of excess non-labeled WT and mutant (M) nucleotide competitors. Arrowheads and asterisks indicate the DNA-protein complexes and free DNAs, respectively. (c) GUS staining of *ProSAUR20:GUS*, *ProSAUR65:GUS*, *ProEXPA5:GUS*, and *ProBGAL2:GUS*. Bars = 1 mm. (d) GUS staining of mutated promoters of *ProSAUR20m:GUS*, *ProSAUR65m:GUS*, *ProEXPA5m:GUS*, and *ProBGAL2m:GUS*. Bars = 1 mm. (e) and (**f**) Reduced GUS activity of *ProSAUR20:GUS tcp_s* seedlings (**e**) and leaves (f). Bars = 1 mm (**e**) and 1 cm (**f**).

### PM H^+^-ATPase activation and apoplast acidification by TCP3

Because SAURs activate PM H^+^-ATPase for cell expansion (Spartz *et al*., 2014; Nagpal *et al*., 2022), we expected that TCP3 would activate PM H^+^-ATPase via induction of the *SAUR20* and *SAUR65* genes. Because phosphorylation of the penultimate threonine residue (Thr_947_) is required for the activation of Arabidopsis H^+^-ATPase2 (AHA2) (Hayashi *et al*., 2010; Takahashi *et al*., 2012) and SAURs mediate Thr_947_ phosphorylation (Spartz *et al*., 2014; Nagpal *et al*., 2022), we compared the Thr_947_ phosphorylation levels of WT, *Pro35S:mTCP3*, and *tcp_s* plants. Immunoblotting analysis indicated that, compared with WT, *Pro35S:mTCP3* exhibited increased Thr_947_ phosphorylation levels, whereas Thr_947_ phosphorylation was decreased in *tcp_s* (Figure **4a** and **4b**, Supporting Information Figure **S10**). These results demonstrate that TCP3 activates PM H^+^-ATPase.

**Figure 4.**
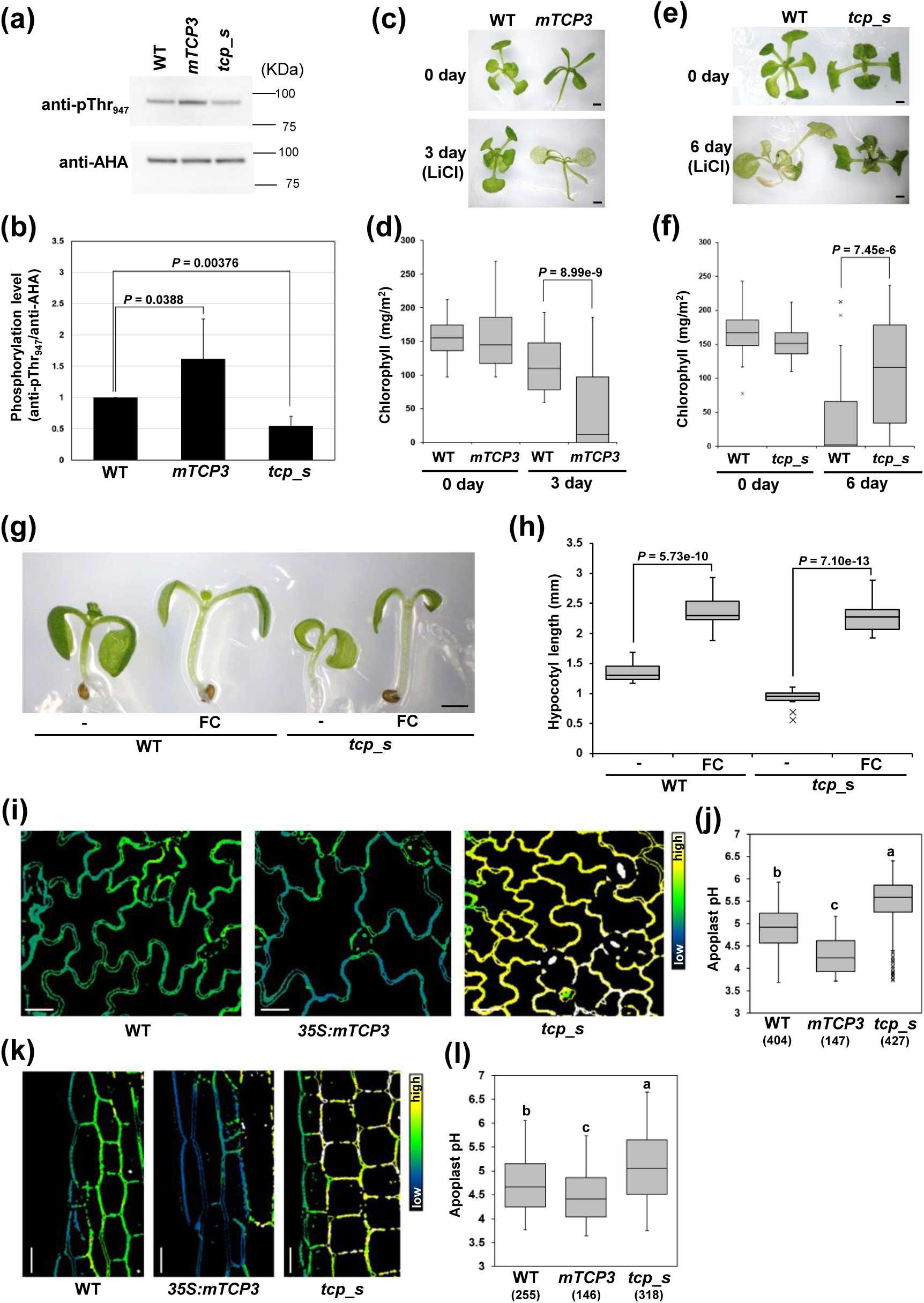
H^+^-ATPase activity and apoplast pH regulated by TCP3. (a) H^+^-ATPase phosphorylation level (pThr_947_) and H^+^-ATPase amount (AHA) were determined by immunoblotting of WT, *Pro35S:mTCP3*, and *tcp_s* using specific antibodies. Molecular size markers are presented to the right of the blots. (b) Quantification of H^+^-ATPase phosphorylation levels of WT, *Pro35S:mTCP3*, and *tcp_s*. H^+^-ATPase phosphorylation levels were quantified as the ratio of the signal intensity of pThr_947_ to that of H^+^-ATPase. Values are the means of six independent experiments, and values for the WT were set at 1. Error bars indicate standard deviation. *P* values were calculated using Dunnett’s test. (c) through (**f**) Images of WT and *Pro35S:mTCP3* (**c**) and WT and *tcp_s* (**e**) grown on plates with or without 20 mM LiCl for the indicated number of days. Bars = 1 mm. Quantification of chlorophyll content in WT and *Pro35S:mTCP3* (**d**) and WT and *tcp_s* (f) grown on plates with 20 mM LiCl. Fifty plants per genotype were analyzed, and the chlorophyll content is shown in the box plots. *P* values were calculated using the Kolmogorov-Smirnov test. Bars = 1mm. (g) Image of WT and *tcp_s* grown with ethanol (−) or fusicoccin (FC). Bar = 1 mm. (h) Hypocotyl length of WT and *tcp_s* grown with ethanol (−) or FC. A total of 12 biological replicates were analyzed, and *P* values were calculated using Student’s *t*-test. (i) through (**l**) HPTS staining of pavement cells of the adaxial cotyledons (**i**) and hypocotyl cells (**k**) of WT, *Pro35S:mTCP3*, and *tcp_s*. Determination of apoplast pH values of cotyledons (**j**) and hypocotyls (**l**) of WT, *Pro35S:mTCP3*, and *tcp_s*. Different letters above the box plots indicate statistically significant differences determined using the Bonferroni test. Sample numbers are indicated in parentheses.

To validate the role of TCP3 in PM H⁺-ATPase–mediated physiological responses, we conducted LiCl and FC assays. The *aha2* mutant reduces the protonmotive force and uptake of toxic cations such as lithium (Haruta *et al*., 2010). We therefore investigated the effect of LiCl on the growth of WT, *Pro35S:mTCP3*, and *tcp_s* plants. Compared with the WT, *Pro35S:mTCP3* was susceptible to LiCl, whereas *tcp_s* was resistant (Figure **4c** to **4f**). Furthermore, FC activates PM H^+^-ATPase, leading to cell expansion (Johansson *et al*., 1993; Spartz *et al*., 2014). We observed that FC enhanced *tcp_s* hypocotyl elongation to a similar extent as WT (Figure **4g** and **4h**). These results were consistent with findings regarding TCP3-activated PM H^+^-ATPase. However, FC did not enhance *Pro35S:mTCP3* hypocotyl elongation, suggesting that *Pro35S:mTCP3* plants activate PM H^+^-ATPase in the absence of FC (Supporting Information Figure **S11a** and **S11b**). In contrast, high temperature induced hypocotyl elongation in the WT but not *tcp_s*, demonstrating the specific effect of FC in enhancing *tcp_s* hypocotyl elongation (Supporting Information Figure **S12a** and **S12b**) (Han *et al*., 2019; Zhou *et al*., 2019).

Because TCP3 activated PM H^+^-ATPase, we expected that TCP3 would induce apoplast acidification. To monitor the apoplast pH dynamics at cellular resolution, we applied the membrane-impermeable dye HPTS as a ratiometric fluorescent pH indicator. HPTS has been used to investigate apoplast pH in Arabidopsis root, hypocotyls, and cotyledon pavement cells (Barbez et al., 2017). HPTS staining indicated that, compared with WT, *Pro35S:mTCP3* ratiometric values were reduced (488/405) in epidermal cotyledon cells and hypocotyl cells, whereas the ratiometric values in *tcp_s* were increased (488/405) (Figure **4i** and **4k**). To interpret these ratiometric changes quantitatively, we converted fluorescence ratios into absolute pH values using the calibration equations (Supporting Information Figure **S13**). This analysis revealed significant differences in apoplast pH among WT, *Pro35S:TCP3*, and *tcp_s* in both cotyledons and hypocotyls (Figures **4j** and **4l**). These results indicate that TCP3 induces apoplast acidification, which could promote cell expansion. In line with the results of extensibility and breaking load of hypocotyls (Figure **1i** and **1j**), the periclinal cell walls of *Pro35S:mTCP3* showed reduced pH compared with those of WT, whereas *tcp_s* showed increased pH (Supporting Information Figure **S14**).

### Genetic interactions between TCP and SAUR genes

To test whether TCPs function upstream of *SAUR* genes *in planta*, we introduced the *gTCP3:GFP-SAUR20* gene into *tcp_s* (*gTCP3:GFP-SAUR20 tcp_s*) and examined whether this could rescue the mutant phenotype. Ectopic expression of *SAUR20* in *tcp_s* substantially recovered the cell volume and hypocotyl length of two independent lines of *gTCP3:GFP-SAUR20 tcp_s* to an extent similar to that of WT plants (Figure **5a** to **5d**). These results indicate that *TCPs* act upstream of *SAUR20*.

**Figure 5.**
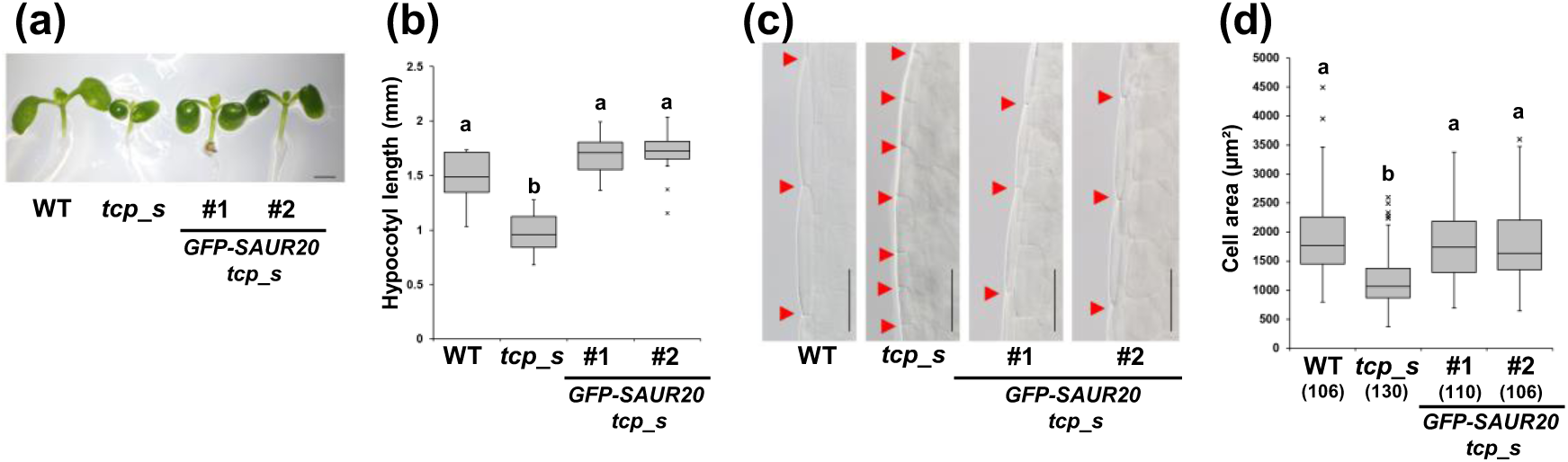
Ectopic expression of *SAUR20* complemented the phenotype of *tcp_s*. (a) Recovery of hypocotyl morphology of WT, *tcp_s*, and two independent *gTCP3:GFP-SAUR20 tcp_s* lines. Bar = 1 mm. (b) Hypocotyl length of WT, *tcp_s*, and two independent *gTCP3:GFP-SAUR20 tcp_s* lines. Different letters above box plots indicate statistically significant differences determined using the Tukey test (n = 12). (c) Hypocotyl cells of WT and two independent *gTCP3:GFP-SAUR20 tcp_s* lines. Scale bars = 100 µm. (d) Size of hypocotyl cells of WT and two independent *gTCP3:GFP-SAUR20 tcp_s* lines. Sample numbers are indicated in parentheses. Different letters above the box plots indicate statistically significant differences determined using the Tukey-Kramer test.

## Discussion

In this study, we focused on the functions of TCPs (Figure **6**), important TFs involved in the morphogenesis of cotyledons, hypocotyls, leaves, and floral organs (Palatnik *et al*., 2003; Koyama *et al*., 2007, 2011; Efroni *et al*., 2008; Wei *et al*., 2015; Huang & Irish, 2015; Challa *et al*., 2016, 2021; Zheng *et al*., 2022; Lan *et al*., 2023). We showed that TCPs promote cell expansion and decrease cell stiffness. TCPs induce the expression of *EXPA5* and *BGAL2* genes encoding cell wall loosening proteins and *SAUR* genes, which activate PM H^+^-ATPase and induce apoplast acidification. The elevated PM H^+^-ATPase activity hyperpolarizes the PM, increases the energy available for solute uptake, maintains water uptake, and thereby allows turgor pressure to induce cell expansion (Du *et al*., 2020). Apoplast acidification activates cell wall loosening proteins (Arsuffi & Braybrook, 2018; Du *et al*., 2020). Acidic pH also protonates auxin and facilitates its diffusion across the PM into cells to promote cell expansion (Arsuffi & Braybrook, 2018). Therefore, TCPs function as master regulators of cell expansion. Importantly, the estimated apoplast pH values (Figures **4j** and **4l**) fell within the range known to activate expansin activity (approximately pH 3.5–5.5; McQueen-Mason et al., 1992, Plant Cell 4:1425–1433), supporting the idea that the observed acidification is biologically meaningful.

**Figure 6.**
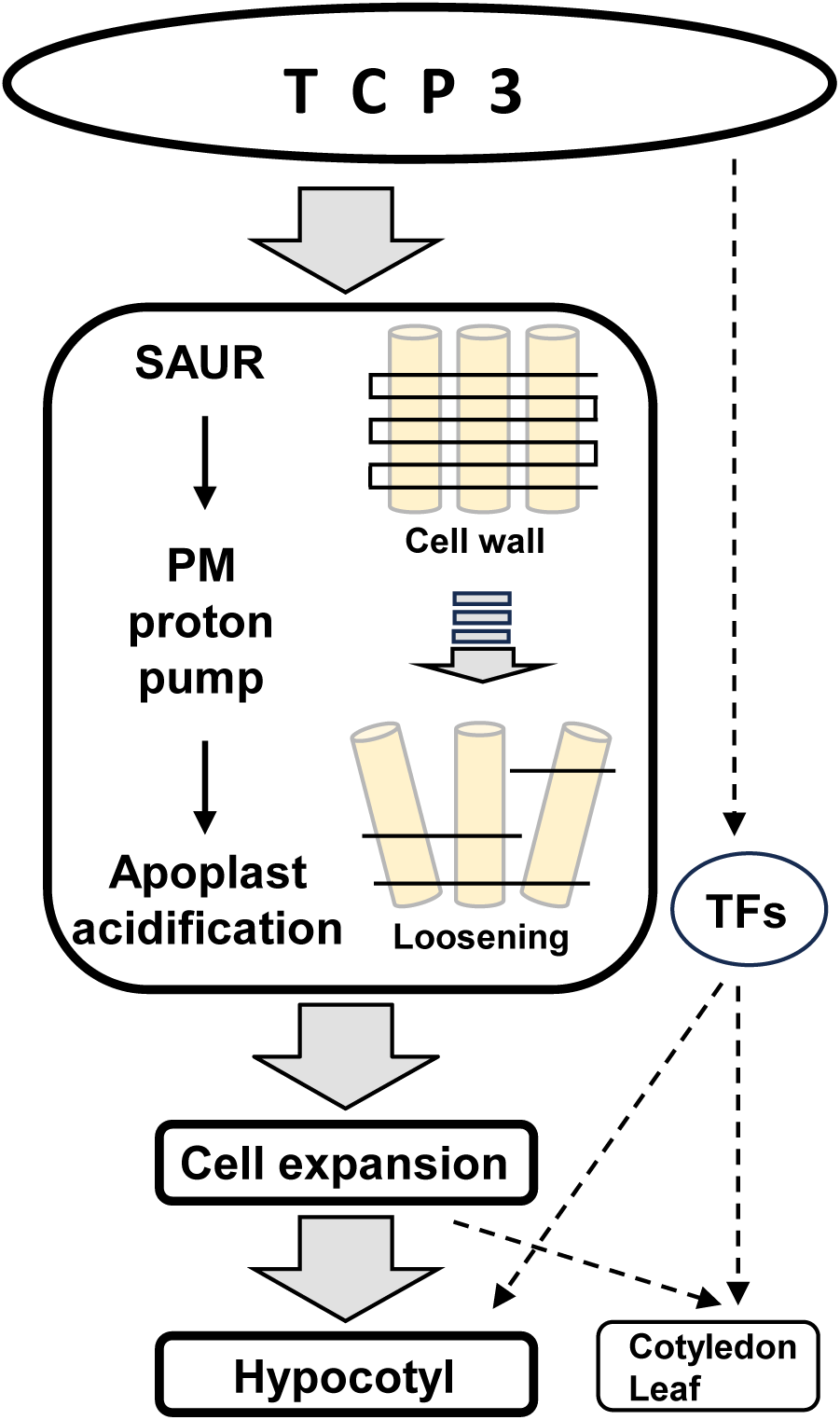
A model of TCP3-mediated regulation of cell expansion. TCP3 promotes apoplast acidification and cell expansion for hypocotyl elongation. In cooperation with downstream transcription factors (TFs), TCP3 controls hypocotyl elongation and development of cotyledons and leaves (dotted arrows). See details in the main text.

Auxin is a well-known inducer of acid growth for cell expansion (Arsuffi & Braybrook, 2018; Du *et al*., 2020). By contrast, the current model for cell expansion is based on direct TCP-mediated activation of the expression of *EXPA5*, *BGAL2,* and *SAUR* genes without exogenous application of auxin. In addition to the auxin-inducible acid growth model, we propose a TCP-promoted cell expansion model. TCP and auxin signals may converge at the level of *SAUR* regulation. In addition to direct regulation of the *SAUR20* and *SAUR65* genes, TCPs regulate auxin signaling pathway genes (Figure **2c**), and auxin induces the expression of *SAUR* genes (Ren & Gray, 2015; Challa *et al*., 2016). Because TCPs interact with auxin-inducible TFs such as ARFs and IAAs (Shani *et al*., 2017; Trigg *et al*., 2017; Altmann *et al*., 2020), TCPs are thought to integrate auxin signals with cell expansion through these protein-protein interactions.

Our data indicate that TCPs directly regulate the expression of *EXPA5*, *BGAL2,* and *SAUR* genes. Dysregulated expression of such genes often induces ectopic cell expansion along with the development of aberrant organ morphologies (Marowa *et al*., 2016). Ectopic micro-induction of expansin genes in leaf margins and shoot apical meristems generates a lobed leaf morphology and irregular phyllotaxis, respectively (Pien et al. 2001). Overexpression of expansin genes leads to larger leaves than WT plants, whereas repression of expansin gene expression results in the development of smaller, wavy leaves (Cho & Cosgrove 2000). Ectopic expression of *SAUR* genes leads to exaggerated elongation of hypocotyls and enlarged leaves (Spartz *et al*., 2014; Vanhaeren *et al*., 2014; Nagpal *et al*., 2022). Our results highlight the critical importance of TCP-mediated regulation of cell expansion in organ morphogenesis. TCPs direct the expression of *SAURs*, *EXPA5*, and *BGAL2* by binding to GGnCCC sequences (Figure **3a**). Our results are in agreement with those of previous reports indicating the requirement of the GGTCCCAT sequence, the core of which consists of the TCP-binding sequence, for the expression of *SAUR* genes (Li *et al*., 1994; Xu *et al*., 1997).

In addition to the direct regulation of *EXPA5*, *BGAL2,* and *SAUR* genes by TCPs as discussed above, we hypothesize that TCPs are involved in the indirect regulation of cell expansion via CUP SHAPED COTYLEDONs (CUCs). CUCs form sinuses in the leaf margins and along cotyledon boundaries (Aida *et al*., 1997; Hasson *et al*., 2011). In contrast to TCPs, CUCs repress the transcription of genes encoding cell wall loosening proteins, resulting in increased cell stiffness and suppression of cell expansion (Bouré *et al*., 2022). Consistent with the repressive regulation of CUCs by TCPs (Koyama *et al*., 2007, 2017; Rubio-Somoza *et al*., 2014), *Pro35S:mTCP3* plants form smooth leaf margins and exhibit cotyledon fusion, which promotes cell expansion (Koyama *et al*., 2007, 2010, 2017). Conversely, the leaf margins of *tcp_s* are serrated, with suppressed cell expansion. The *CUC*-mediated pathways may augment TCP-mediated regulation of cell expansion.

TCPs also induce the expression of *TF* genes required for hypocotyl elongation (Figure **2c**, Figure **6**). Furthermore, TCPs regulate plant hormone signaling pathways (Koyama *et al*., 2010; Efroni *et al*., 2013; Challa *et al*., 2016, 2019) and interact with other TFs to modulate the extent of cell expansion (Alvarez *et al*., 2016; Tao *et al*., 2013). Of particular importance, TCPs regulate the expression of *SAUR16* and *SAUR50*, which are categorized differently than *SAUR20* and *SAUR65* subfamily genes, and this regulation promotes seedling de-etiolation processes, such as cotyledon opening (Dong *et al*., 2019). The sequential regulation of different sets of *SAUR* genes by TCPs affects a broad range of processes, from de-etiolation to light-grown seedling morphogenesis. In this way, TCPs integrate additional signals to fine-tune cell expansion. On the other hand, ectopic overexpression of *SAUR20* did not complement the cotyledons and leaves of *tcp_s* (Figure **5**), suggesting that additional factors downstream of TCPs might be required to develop these organs (Figure **6**).

The results of our study provide new insights into the regulation of cell expansion during leaf morphogenesis. During leaf morphogenesis, TCPs promote the transition from cell division to cell expansion, which follows a basipetal gradient (Nath *et al*., 2003; Efroni *et al*., 2008; Bresso *et al*., 2018; Tsukaya, 2021). The delayed transition observed in *tcp* mutant leaves is believed to induce mechanical stress, which results in a buckled leaf morphology (Trinh *et al*., 2021). TCPs suppress the activity of cyclin to halt cell division (Efroni *et al*., 2008; Schommer *et al*., 2014; Bresso *et al*., 2018), but the mechanism of the TCP-promoted transition to cell expansion remains unknown. We found no clear evidence of a direct connection between TCP3 and cell division (Figure 2C). As shown in Supporting Information Figure **S15**, *ProSAUR20:GUS* signaling exhibited a basipetal gradient that contrasted with the pattern of cell division markers (Donnelly *et al*., 1999; Challa *et al*., 2019). This suggests that *SAUR20* mediates the basipetal transition to cell expansion. The functionally redundant TCPs finely regulate their own expression domains, which follow the basipetal gradient, through a double negative feedback loop between the TCPs and miR319 (Shankar et al. 2023). Based on this, we speculate that these TCPs precisely specify the spatial and temporal boundary of cell division and expansion. Our results support the important contributions of TCP-promoted cell expansion to leaf morphogenesis.

Finally, we note important considerations regarding our mechanical and pH measurements. In hypocotyls, tensile testing showed that TCP3 induced cell wall loosening (Figure **1i** and **1j**), and HPTS staining indicated that TCP3 lowered the pH of the periclinal cell walls (Supplemental Figure **S14**). Both findings support our model of TCP3-mediated cell expansion. In cotyledons, however, mechanical and pH measurements were performed on different wall orientations. While AFM probes contacted periclinal walls, HPTS assessed the pH of anticlinal walls. Because AFM measurements on anticlinal walls are technically challenging (Supplemental Figure **S16**; Peaucelle et al., 2015; Bidhendi and Geitmann, 2019), we could not measure stiffness on the same wall face where pH was measured in cotyledons. This discrepancy in measurement sites should be taken into account when interpreting the cotyledon data. Furthermore, our AFM experiments in cotyledons were performed under both turgid and plasmolyzed conditions (Figure **1h**; Supplemental Figure **S16**). Measurements on turgid cells can be influenced by internal turgor pressure, whereas measurements on plasmolyzed cells remove turgor effects but may introduce artifacts such as altered wall polymer hydration, loss of plasma membrane support, and cell deformation (Braybrook, 2015; Bidhendi and Geitmann, 2019). These context-dependent effects could explain discrepancies in the Young’s modulus values obtained under turgid versus plasmolyzed conditions. A more cautious and accurate interpretation is possible when these technical differences and potential artifacts are taken into account.

## Acknowledgements

We thank Dr. Nam Hai Chua for providing *pER8* plasmids, Drs. Tomohiro Osugi and Yuta Takase for helpful discussions, and Ms. Eriko Tanaka for skilled technical assistance. This work was supported by KAKENHI JP25K09671 to T.Koyama, JP24H02252 to T.Kunieda, and JP18H05484 to T.D.

## Author Contributions

T.Koyama designed the research; T.Koyama., T.Kunieda., H.T., M.N., K.S., N.M., J.I., M.S., A.B., and T.D. performed the research; K.T., T.Kinoshita., and M.O.-T. contributed new reagents/analytical tools; T.Koyama., T.Kunida, and .N.M. wrote the paper.

## Data availability

The data that support the findings of this study are available in the main text and Supporting Information Figures S1–S16 and Tables S1–S2 of this article.

## Supporting Information for

### This Supporting Information includes

Supporting Information Figures **S1** to **S16**

Supporting Information Table **S1 to S2** (Table **S2** is provided as a separate Excel file)

**Supporting Information Figure S1.**
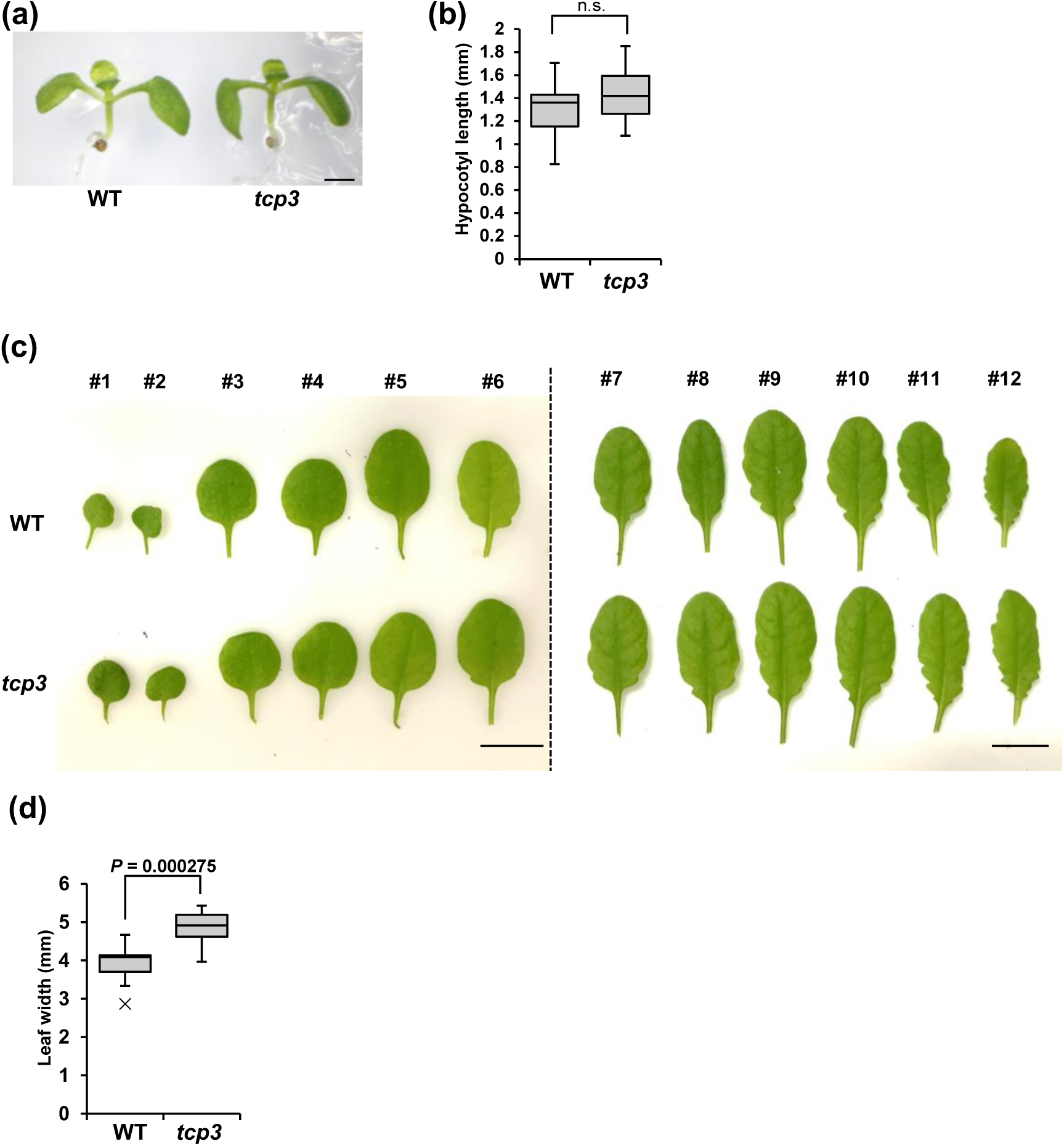
Morphology of WT and *tcp3* single mutant. **(a)** Morphology of 7-day-old WT and *tcp3* mutant seedlings. Bar = 1 mm. **(b)** Hypocotyl length was not significantly different between WT and *tcp3* mutant seedlings. “n.s.” indicates no significant difference based on Student’s t-test (n = 12). **(c)** Comparison of leaf morphology between WT and *tcp3* mutant. The first to sixth leaves were detached from 4-week-old plants, and the seventh to twelfth leaves from 5-week-old plants. The overall leaf shape and margins were comparable between WT and *tcp3* mutant. Bars = 1 cm. **(d)** Leaf width of the first and second leaves in WT and *tcp3* mutant. Statistically significant differences were determined using Student’s t-test (n = 12). The increased leaf width observed in *tcp3* is similar to phenotypes reported for *tcp4* and *tcp10* single mutants, and may represent a mild effect of *TCP* gene disruption (Schommer et al., 2008).

**Supporting Information Figure S2.**
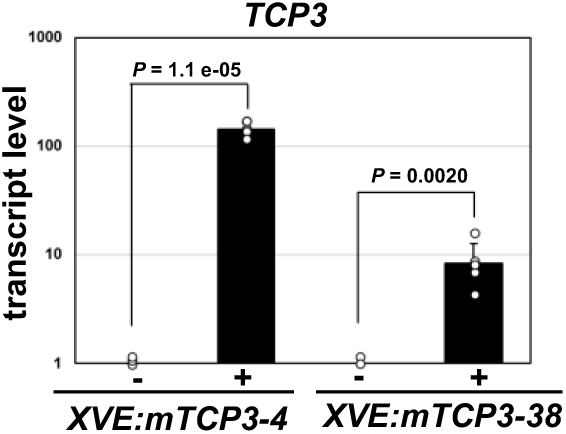
*TCP3* transcript levels in *ProXVE:mTCP3* incubated with DMSO (−) or estradiol (+) for 16 h. Error bars indicate standard deviations of six biological replicates, and *P* values were calculated using Student’s *t*-test.

**Supporting Information Figure S3.**
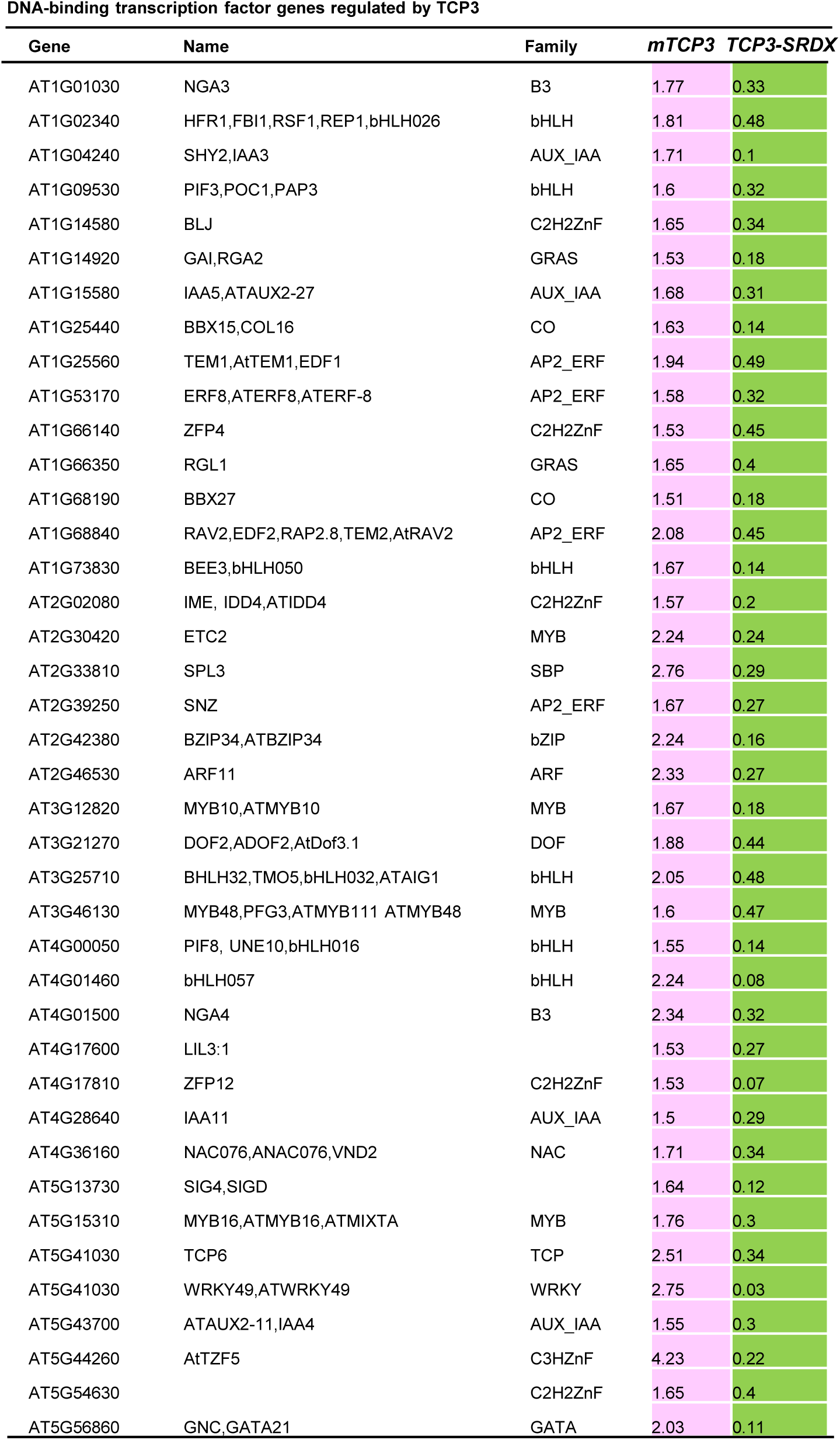
Expression of DNA-binding TF genes in *ProXVE:mTCP3* and *ProXVE:TCP3-SRDX* plants.

**Supporting Information Figure S4.**
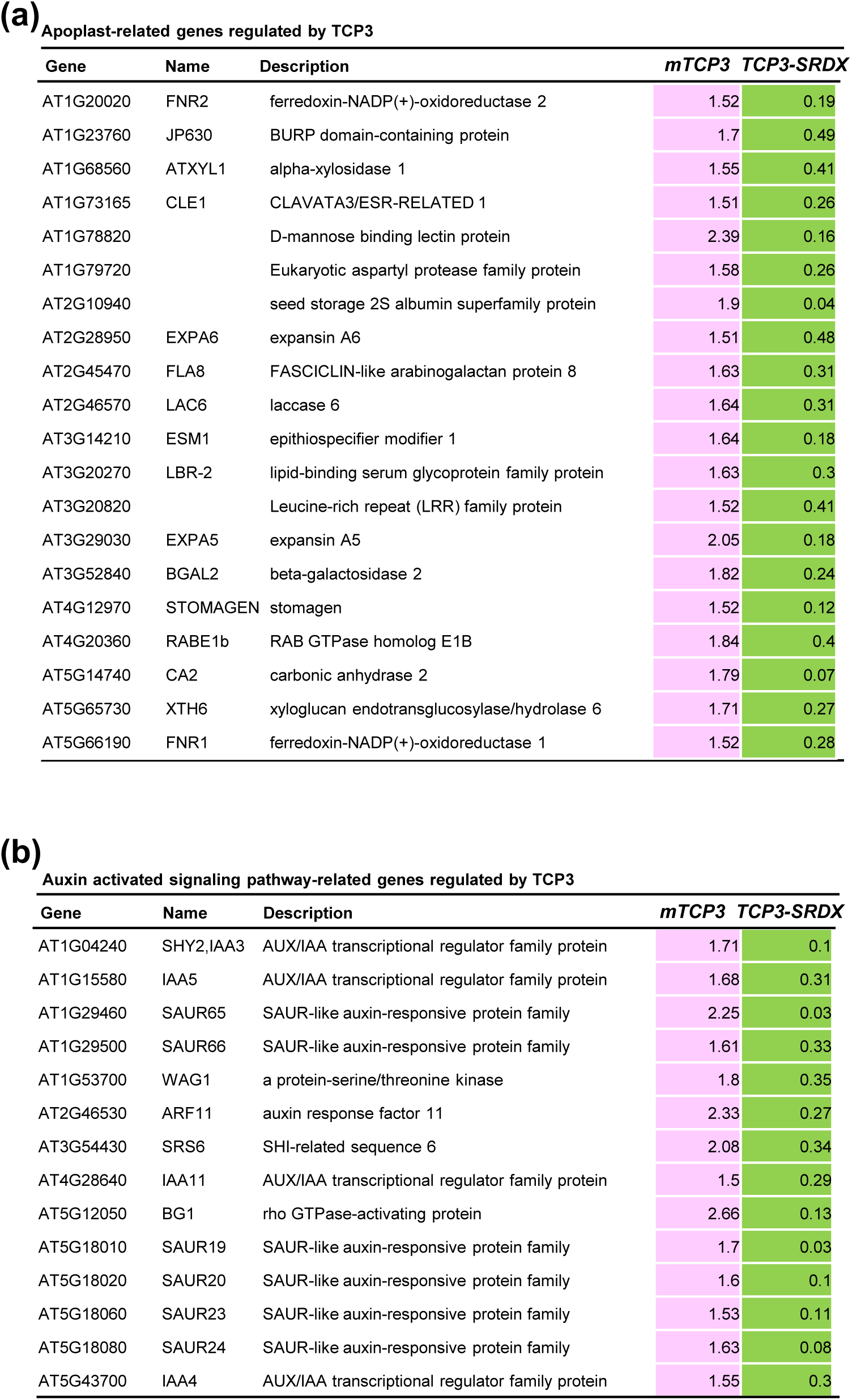
Apoplast-related and auxin-activated signaling pathway–related genes downstream of *TCP3*. **(a)** Expression of apoplast-related genes in *ProXVE:mTCP3* and *ProXVE:TCP3-SRDX* plants. **(b)** Expression of auxin-activated signaling pathway–related genes in *ProXVE:mTCP3* and *ProXVE:TCP3-SRDX* plants.

**Supporting Information Figure S5.**
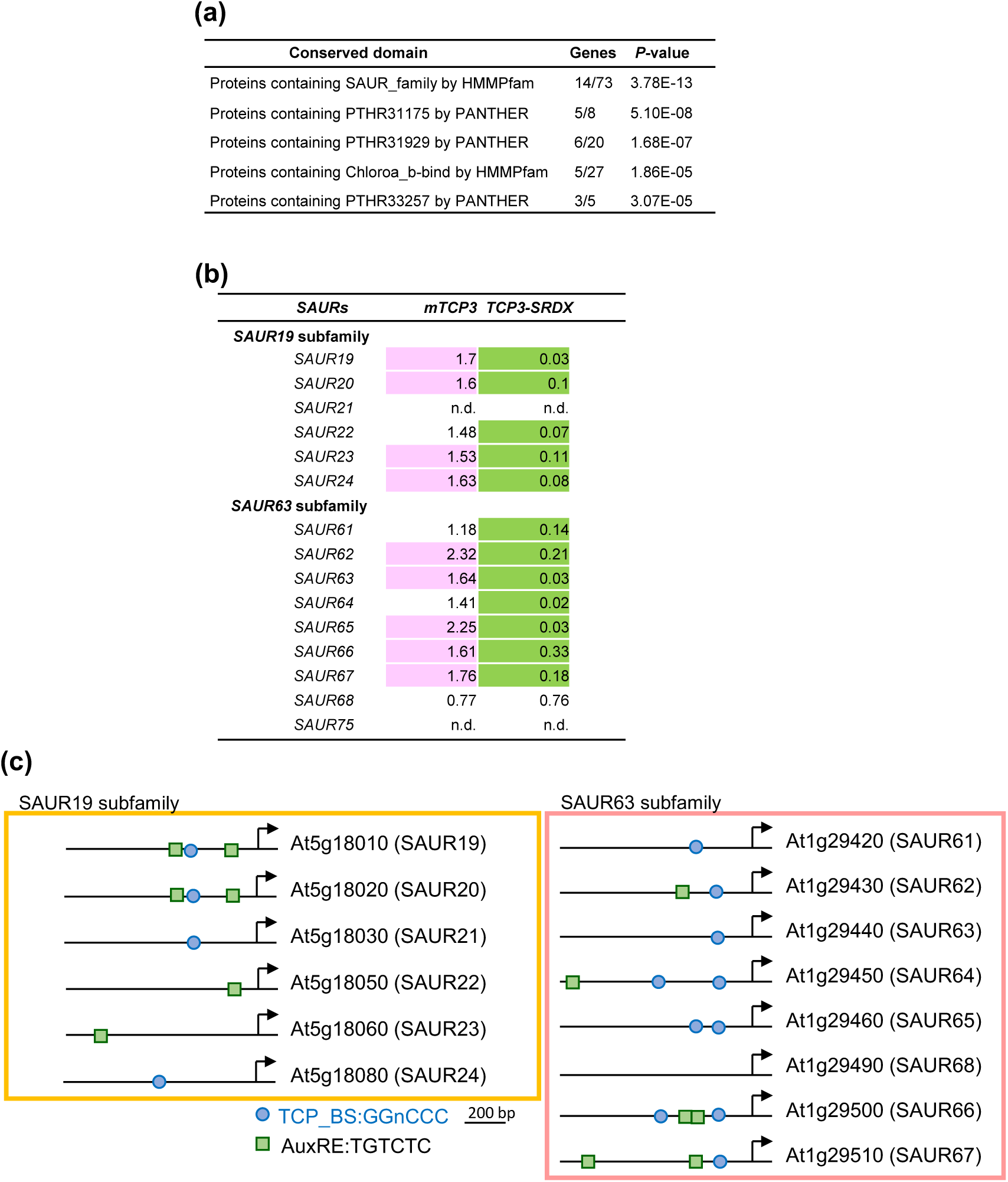
*SAUR19* and *SAUR63* subfamilies downstream of *TCP3*. **(a)** A group of the conserved domain of *SAUR* family enriched in *TCP3* downstream genes. **(b)** Expression of *SAUR19* and *SAUR63* subfamilies in *ProXVE:mTCP3* and *ProXVE:TCP3-SRDX* plants. **(c)** Putative TCP-binding sequences (TCP_BS; circles) and auxin-responsive elements (AuxRE; squares) in the promoters of *SAUR19* and *SAUR63* subfamily genes.

**Supporting Information Figure S6.**
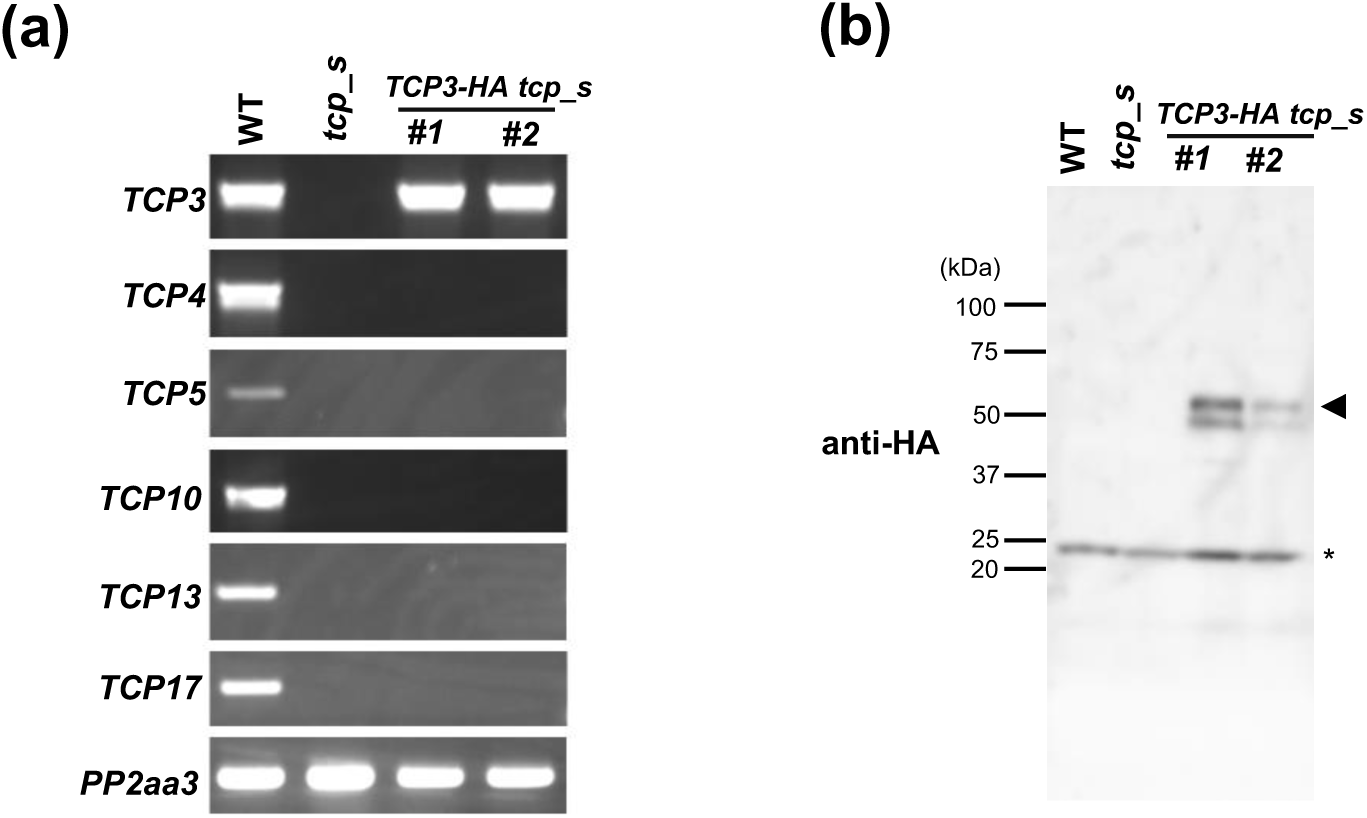
Detection of TCP3-HA in *gTCP3:TCP3-HA tcp_s*. **(a)** RT-PCR analysis of *TCP* transcripts in WT, *tcp_s*, and two independent *gTCP3:TCP3-HA tcp_s* lines. *PP2aa3* genes were analyzed as controls. **(b)** Immunoblotting analysis of WT, *tcp_s*, and two independent *gTCP3:TCP3-HA tcp_s* lines. Immunoreactive bands (arrowhead) were observed in the two *gTCP3:TCP3-HA tcp_s* lines, but not WT or *tcp_s*, demonstrating the accumulation of TCP3-HA. Asterisk indicates nonspecific bands that served as an internal control. Molecular weight markers are shown at the left of the blot.

**Supporting Information Figure S7.**
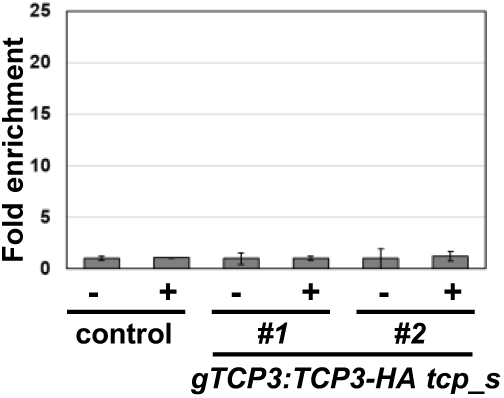
ChIP analysis of *UBQ10*. Chromatin from *tcp_s* (control) and *gTCP3:TCP3-HA tcp_s* was immunoprecipitated with (+) or without (−) anti-HA antibody, and the precipitated DNA fragments were quantified using real-time PCR. Fold-enrichment was determined relative to the value calculated in the absence of antibodies. Error bars indicate the standard deviation of technical triplicates.

**Supporting Information Figure S8.**
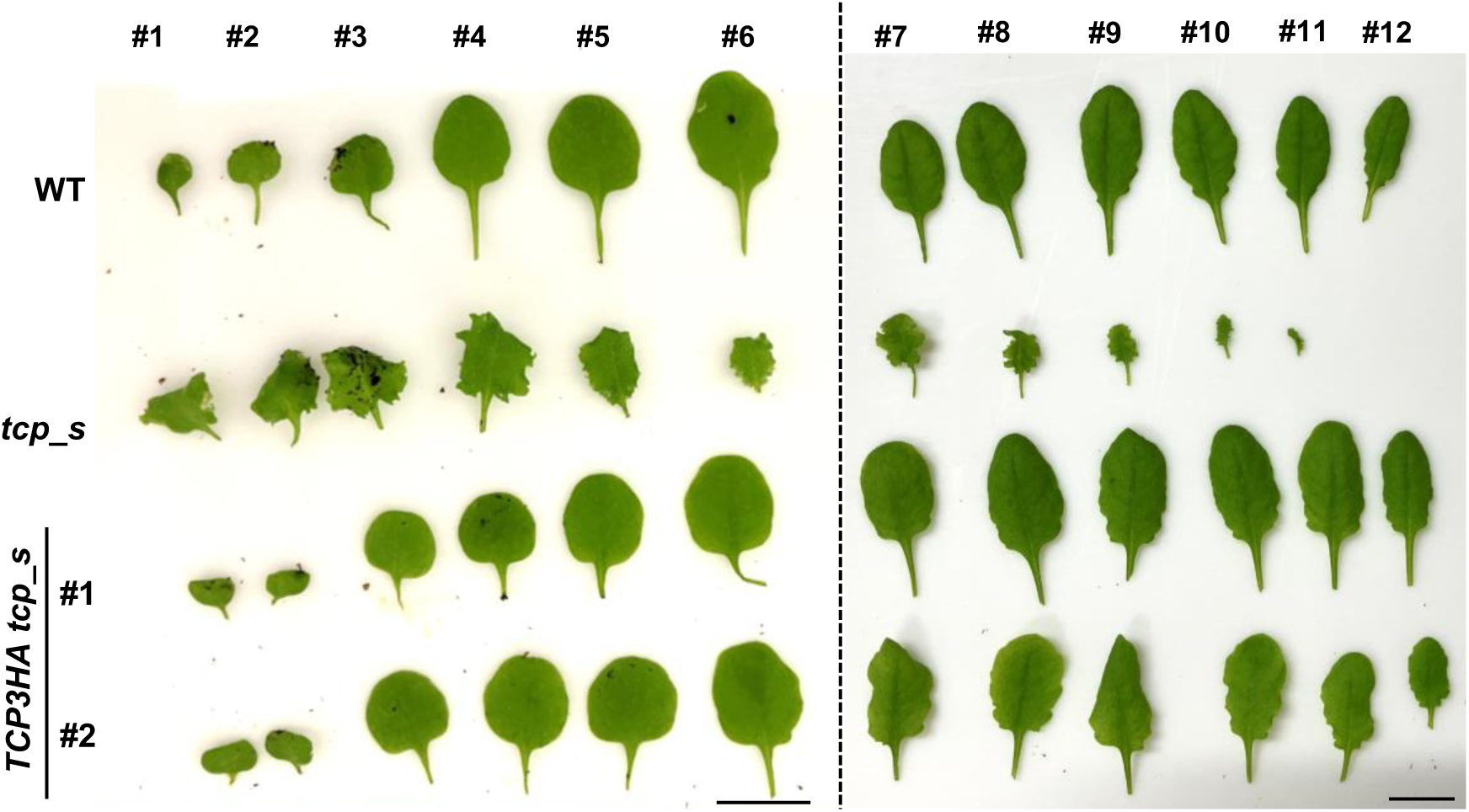
Leaf morphology of WT, *tcp_s* mutant, and two independent lines of *gTCP3:TCP3-HA tcp_s* plants. The first to sixth and the seventh to twelfth leaves were detached from 4-week-old and 5-week-old rosettes, respectively. The first to sixth leaves of *gTCP3:TCP3-HA tcp_s* plants exhibited normal morphology, whereas the seventh to twelfth leaves showed mild waviness and serration. The twelfth leaf was absent in *tcp_s* due to a longer plastochron length. Bars = 1 cm.

**Supporting Information Figure S9.**
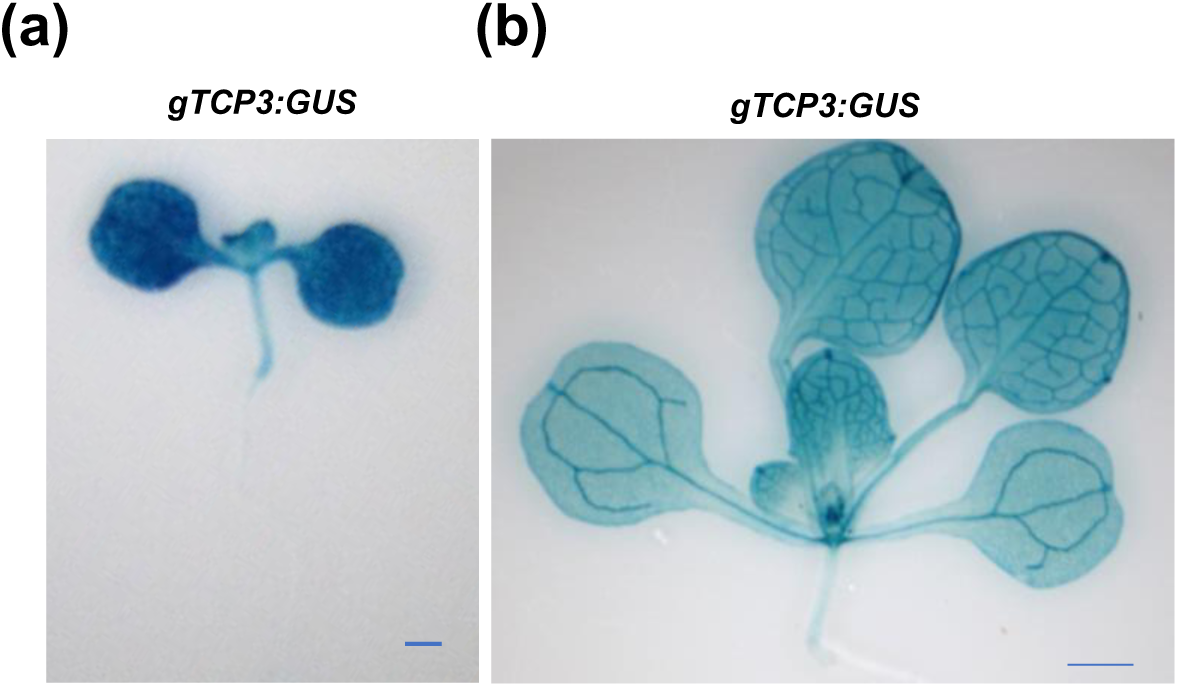
GUS staining of *gTCP3:GUS* seedling **(a)** and rosette **(b)**. Bars = 1 mm.

**Supporting Information Figure S10.**
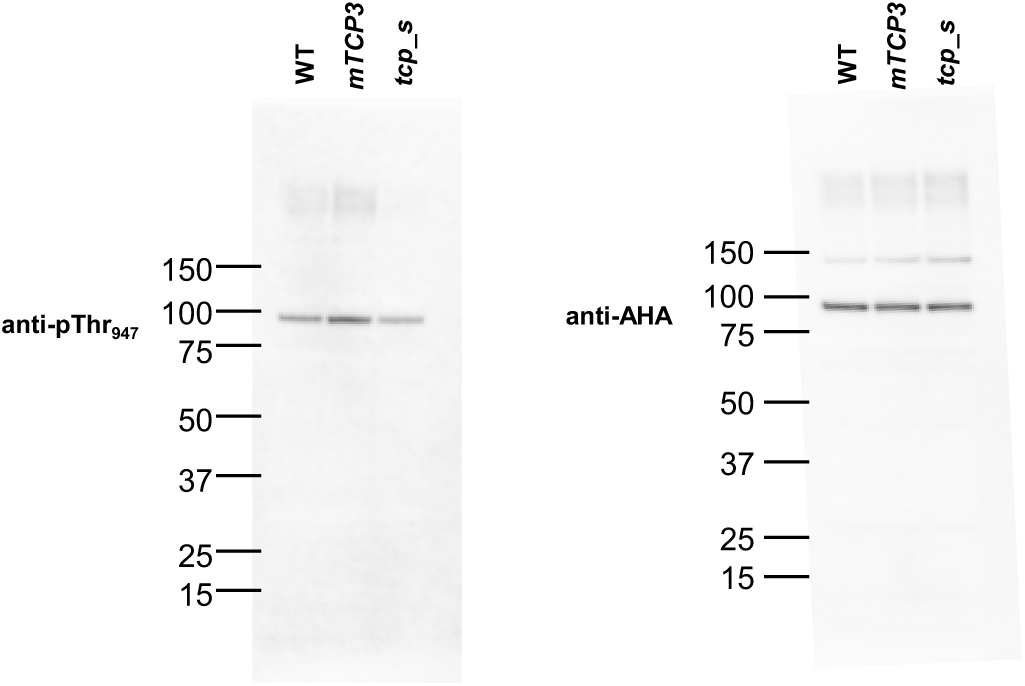
Immunoblotting analysis of WT, *Pro35S:mTCP3*, and *tcp_s* using anti-pTHr_947_ and anti-AHA specific antibodies. Enlarged views of the active signals are shown in Figure 4 **(a)**. Molecular weight markers (KDa) are presented at the left of the blots.

**Supporting Information Figure S11.**
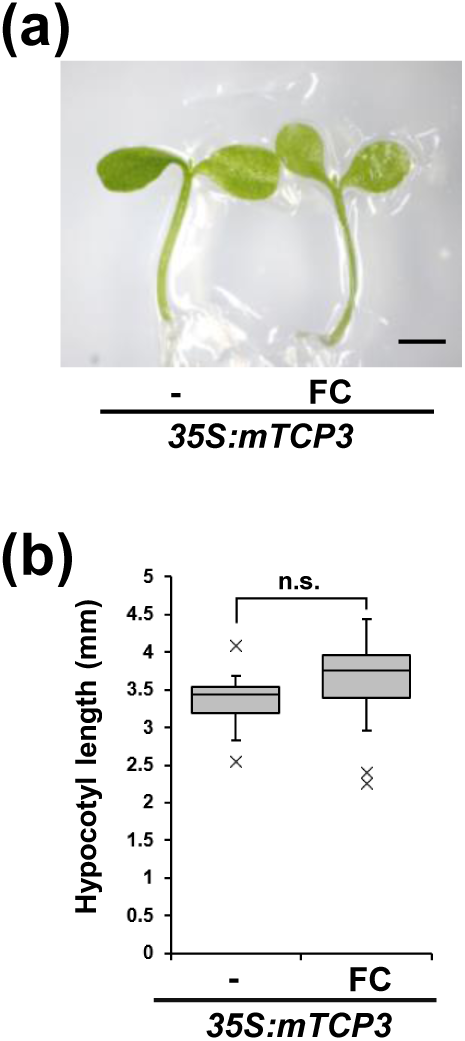
*Pro35S:mTCP3* did not respond to FC. **(a)** Image of *Pro35S:mTCP3* grown with ethanol (−) or FC. Bar = 1 mm. **(b)** Hypocotyl length of *Pro35S:mTCP3* grown with ethanol (−) or FC. A total of 12 biological replicates were analyzed, and *P* values were calculated using Student’s *t*-test. n.s. presents not significant.

**Supporting Information Figure S12.**
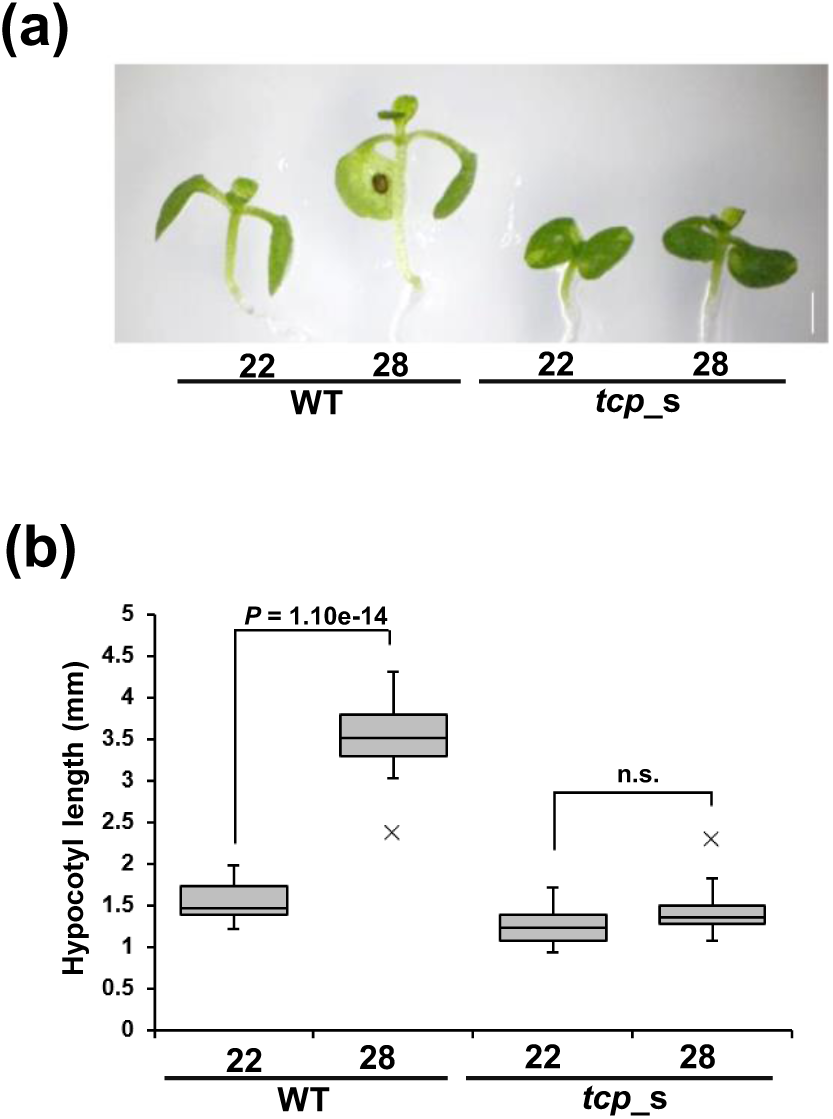
*tcp_s* did not respond to high-temperature stress. **(a)** Image of WT and *tcp_s* grown at 22°C or 28°C for 7 days. Bar = 1 mm. **(b)** Hypocotyl length of WT and *tcp_s* grown at 22°C or 28°C. A total of 15 biological replicates were analyzed, and *P* values were calculated using Student’s *t*-test. n.s. presents not significant.

**Supporting Information Figure S13.**
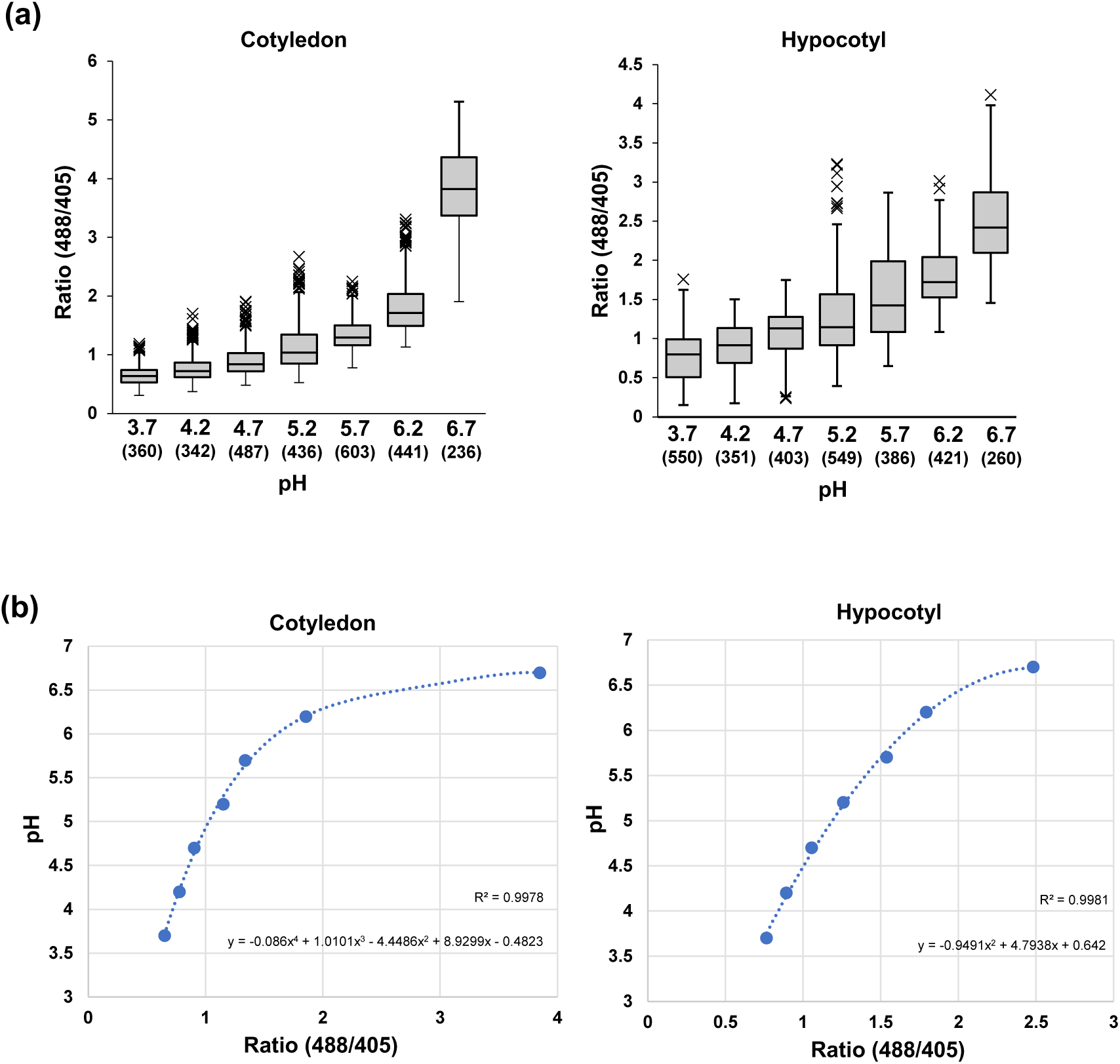
Calibration of HPTS fluorescence ratios (488/405) to absolute pH values. **(a)** WT cotyledons and hypocotyls were incubated in buffer solutions ranging from pH 3.7 to 6.7. Fluorescence ratios (488/405) were measured and presented as box plots. [X-axis: pH; Y-axis: fluorescence ratio (488/405)]. Sample numbers are indicated in parentheses. **(b)** Calibration curves for cotyledons and hypocotyls were generated using the mean ratios from **(a)**. The regression curves were used to estimate pH from fluorescence ratios. [X-axis: fluorescence ratio (488/405); Y-axis: pH]

**Supporting Information Figure S14.**
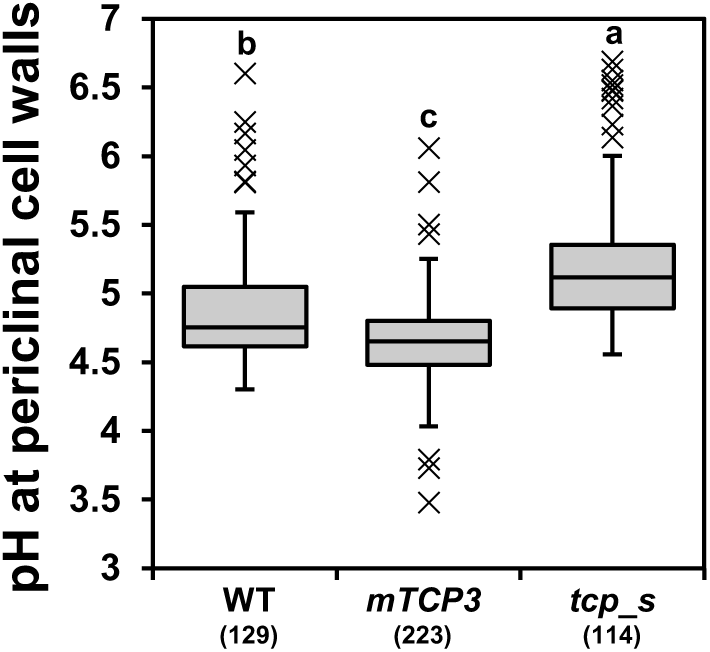
Determination of the pH of the periclinal cell wall in WT, *Pro35S:mTCP3*, and *tcp_s*. Different letters above the box plots indicate statistically significant differences determined using the Bonferroni test. Sample numbers are indicated in parentheses.

**Supporting Information Figure S15.**
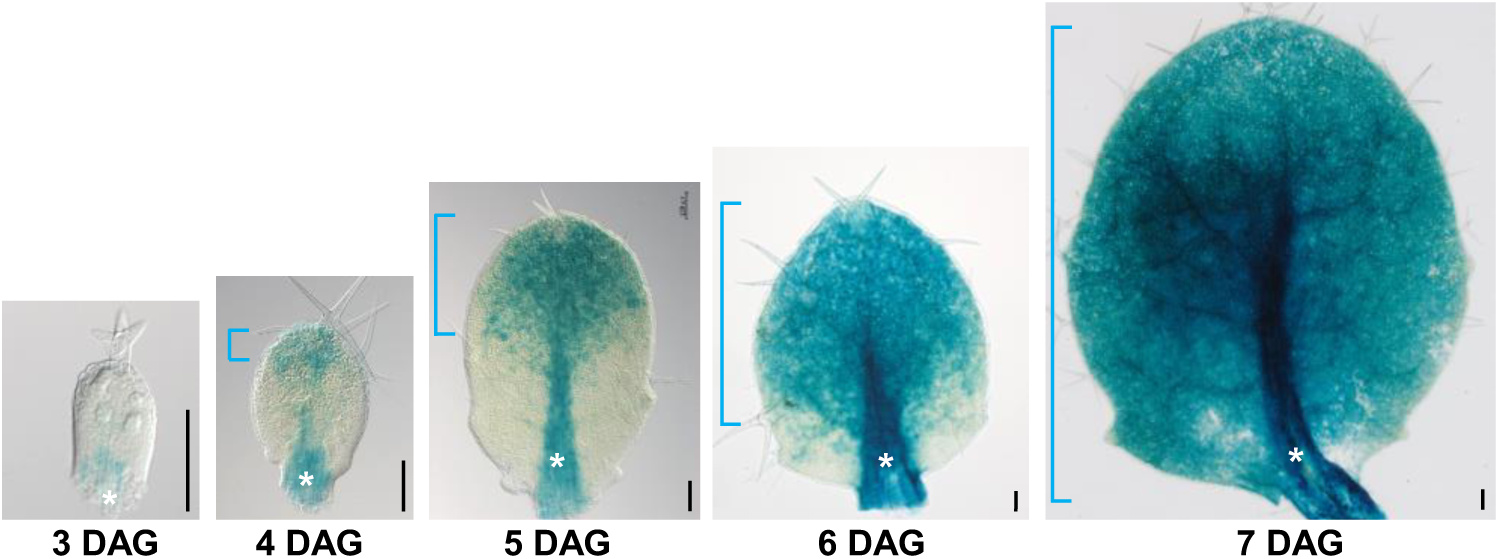
Basipetal gradient of the *ProSAUR20:GUS* signal in leaves. *ProSAUR20:GUS* signals in leaves of seedlings at three to seven days after germination (DAG). Brackets represent the basipetal gradient of *ProSAUR20*-active regions. Asterisks indicate *ProSAUR20* signals that overlap with the midribs and petioles. Bars = 100 µm.

**Supporting Information Figure S16.**
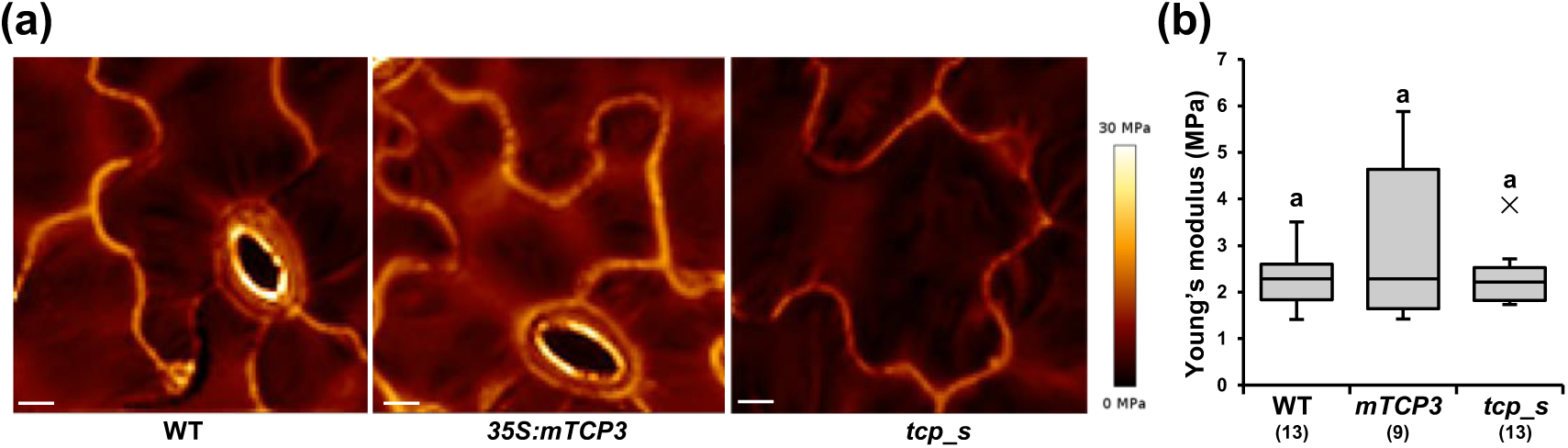
Young’s modulus of WT, *Pro35S:mTCP3* and *tcp_s* in plasmolyzed conditions. (a) AFM images of the adaxial pavement cells of WT, *Pro35S:mTCP3*, and *tcp_s* cotyledons. The color code of Young’s modulus is presented to the right of the images. Scale bars = 10 µm. (b) Quantification of the Young’s modulus in WT, *Pro35S:mTCP3*, and *tcp_s* cotyledon cells under plasmolyzed conditions. Data are presented as box plots. Sample numbers are indicated in parentheses. The letter “a” above all box plots denotes no statistically significant difference, as determined by the Tukey–Kramer test. The tissues were plasmolyzed with 0.55 M mannitol. The measurement protocol followed that used for the turgid condition shown in Figure 1, except that the scan speed was set to 62.5 μm/s. The average indentation depth was approximately 135 nm.

**Supporting information Table S1.**
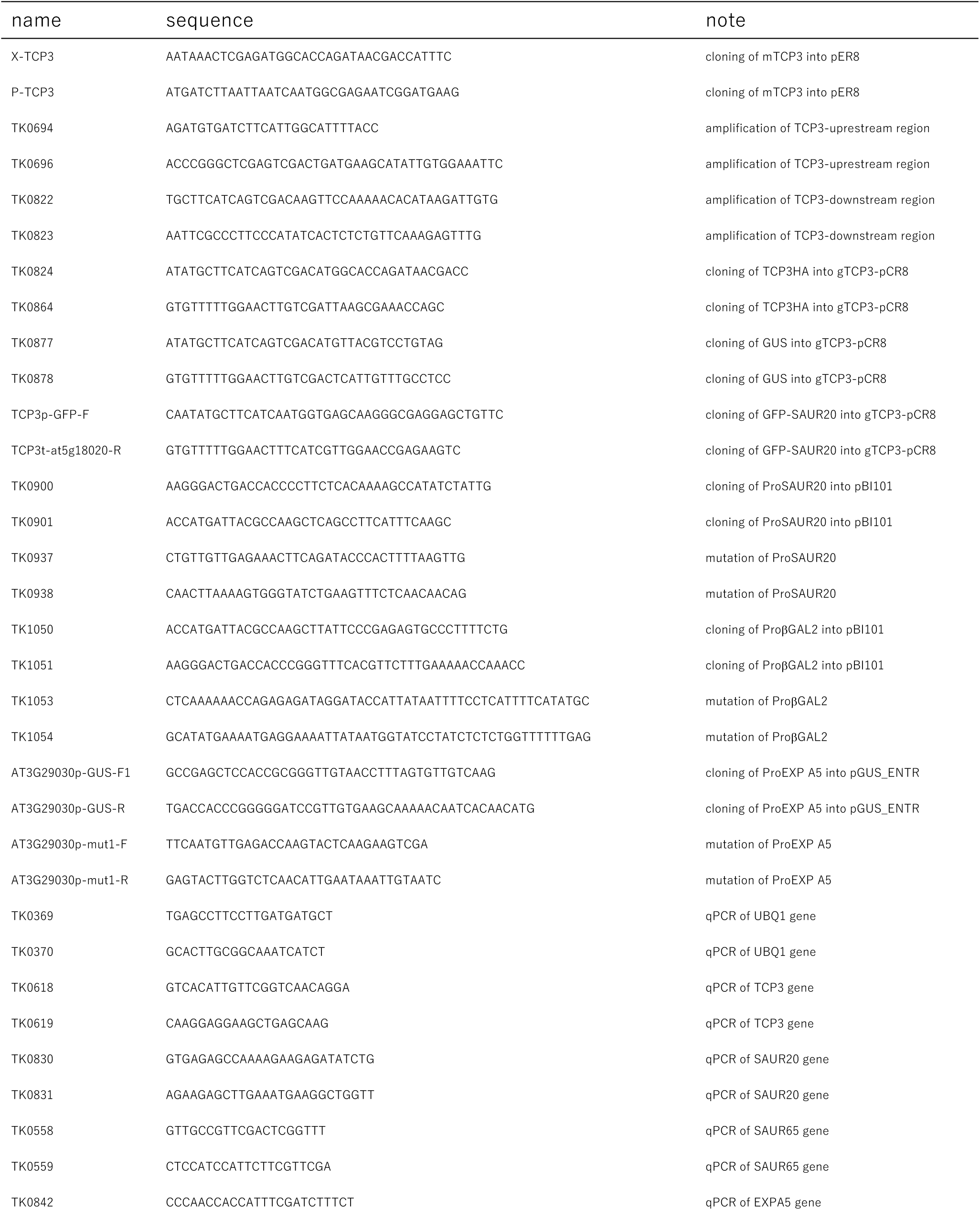
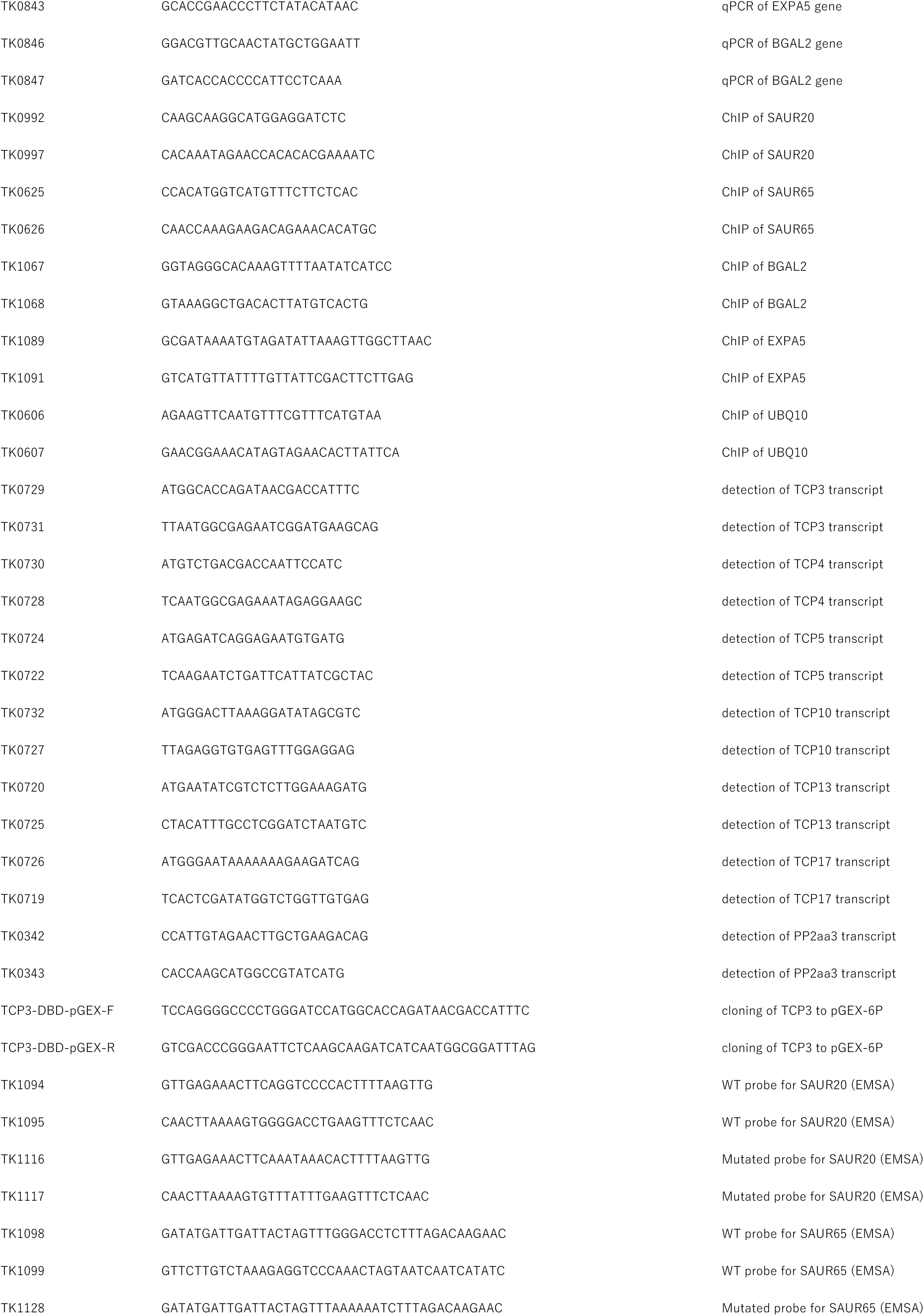
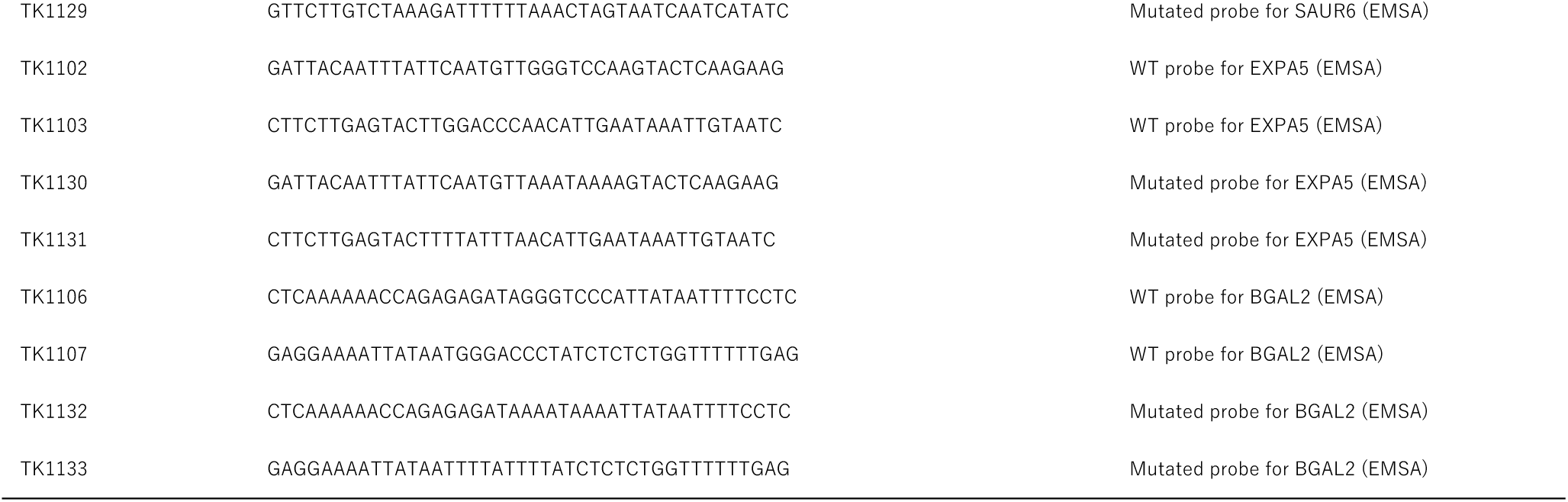
Primers used in this study.

Table **S2** Genes downstream of TCP3.

## References

Aida M, Ishida T, Fukaki H, Fujisawa H, Tasaka M. 1997. Genes involved in organ separation in *Arabidopsis*: An analysis of the cup-shaped cotyledon mutant. Plant Cell 9: 841–857.

Altmann M, Altmann S, Rodriguez PA, Weller B, Elorduy Vergara L, Palme J, Marín-de la Rosa N, Sauer M, Wenig M, Villaécija-Aguilar JA, et al. 2020. Extensive signal integration by the phytohormone protein network. Nature 583: 271–276.

Alvarez JP, Furumizu C, Efroni I, Eshed Y, Bowman JL. 2016. Active suppression of a leaf meristem orchestrates determinate leaf growth. eLife 5: e15023.

Arsuffi G, Braybrook SA. 2018. Acid growth: An ongoing trip. Journal of Experimental Botany 69: 137–146.

Barbez E, Dünser K, Gaidora A, Lendl T, Busch W. 2017. Auxin steers root cell expansion via apoplastic pH regulation in *Arabidopsis thaliana*. Proceedings of National Academy of Science of USA 114: E4884–E4893.

Bidhendi AJ, and Geitmann A. 2019. Methods to quantify primary plant cell wall mechanics. Journal of Experimental Botany 70: 3615–3648.

Boron AK, Vissenberg K. 2014. The *Arabidopsis thaliana* hypocotyl, a model to identify and study control mechanisms of cellular expansion. Plant Cell Reports 33: 697–706.

Bouré N, Peaucelle A, Goussot M, Adroher B, Soubigou-Taconnat L, Borrega N, Biot E, Tariq Z, Martin-Magniette ML, Pautot V, et al. 2022. A cell wall-associated gene network shapes leaf boundary domains. Development 149: dev200359 (2022).

Braybrook SA. 2015 Measuring the elasticity of plant cells with atomic force microscopy. Methods in Cell Biology 125: 237–254.

Bresso EG, Chorostecki U, Rodriguez RE, Palatnik JF, Schommer C. 2018. Spatial control of gene expression by miR319-regulated TCP transcription factors in leaf development. Plant Physiology 176: 1694–1708.

Cendreau E, Traas J, Demos’ T, Crandjean O, Caboche M, Hofte H. 1997. Cellular basis of hypocotyl growth in *Arabidopsis thaliana*. Plant Physiology 114: 295–305.

Challa KR, Aggarwal P, Nath U. 2016. Activation of YUCCA5 by the transcription factor TCP4 integrates developmental and environmental signals to promote hypocotyl elongation in *Arabidopsis*. Plant Cell 28: 2117–2130.

Challa KR, Rath M, Nath U. 2019. The CIN-TCP transcription factors promote commitment to differentiation in Arabidopsis leaf pavement cells via both auxin-dependent and independent pathways. PLoS Genetics 15: e1007988.

Challa KR, Rath M, Sharma AN, Bajpai AK, Davuluri S, Acharya KK, Nath U. 2021. Active suppression of leaflet emergence as a mechanism of simple leaf development. Nature Plants 7: 1264–1275.

Chebli Y, Geitmann A. 2017. Cellular growth in plants requires regulation of cell wall biochemistry. Current Opinion in Cell Biology 44: 28–35.

Cho H-T, Cosgrove DJ. 2000. Altered expression of expansin modulates leaf growth and pedicel abscission in *Arabidopsis thaliana*. Proceedings of National Academy of Science of USA 97: 9783–9788.

Cifuentes-Esquivel N, Bou-Torrent J, Galstyan A, Gallemí M, Sessa G, Salla Martret M, Roig-Villanova I, Ruberti I, Martínez-García JF. 2013. The bHLH proteins BEE and BIM positively modulate the shade avoidance syndrome in *Arabidopsis* seedlings. Plant Journal 75: 989–1002.

Cosgrove DJ. 2023. Structure and growth of plant cell walls. Nature Reviews Molecular Cell Biology 25: 340–358.

Cowling RJ, Harberd NP. 1999. Gibberellins control *Arabidopsis* hypocotyl growth via regulation of cellular elongation. Journal of Experimental Botany 50: 1351–1357.

Dong J, Sun N, Yang J, Deng Z, Lan J, Qin G, He H, Deng XW, Irish VF, Chen H, et al. 2019. The transcription factors TCP4 and PIF3 antagonistically regulate organ-specific light induction of saur genes to modulate cotyledon opening during de-etiolation in *Arabidopsis*. Plant Cell 31: 1155–1170.

Donnelly PM, Bonetta D, Tsukaya H, Dengler RE, Dengler NG. 1999. Cell cycling and cell enlargement in developing leaves of *Arabidopsis*. Developmental Biology 215: 407–419.

Du M, Spalding EP, Gray WM. 2020. Rapid Auxin-Mediated Cell Expansion. Annual Review of Plant Biology 71: 379–402.

Efroni I, Blum E, Goldshmidt A, Eshed Y. 2008. A protracted and dynamic maturation schedule underlies *Arabidopsis* leaf development. Plant Cell 20: 2293–2306.

Efroni I, Han SK, Kim HJ, Wu MF, Steiner E, Birnbaum KD, Hong JC, Eshed Y, Wagner D. 2013. Regulation of leaf maturation by chromatin-mediated modulation of cytokinin responses. Developmental Cell 24: 438–445.

Fairchild CD, Schumaker MA, Quail PH. 2000. HFR1 encodes an atypical bHLH protein that acts in phytochrome A signal transduction. Genes and Development 14: 2377–2391.

Friml J, Gallei M, Gelová Z, Johnson A, Mazur E, Monzer A, Rodriguez L, Roosjen M, Verstraeten I, Živanović BD, et al. 2022. ABP1–TMK auxin perception for global phosphorylation and auxin canalization. Nature 609: 575–581.

Gantulga D, Turan Y, Bevan DR, Esen A. 2008. The Arabidopsis At1g45130 and At3g52840 genes encode β-galactosidases with activity toward cell wall polysaccharides. Phytochemistry 69: 1661–1670.

Geilfus C-M. 2017. The pH of the apoplast: dynamic factor with functional impact under stress. Molecular Plant 10: 1371–1386.

Goh HH, Sloan J, Dorca-Fornell C, Fleming A. 2012. Inducible repression of multiple expansin genes leads to growth suppression during leaf development. Plant Physiology 159: 1759–1770.

Han X, Yu H, Yuan R, Yang Y, An F, Qin G. 2019. *Arabidopsis* transcription factor TCP5 controls plant thermomorphogenesis by positively regulating PIF4 activity. iScience 15: 611–622.

Haruta M, Burch HL, Nelson RB, Barrett-Wilt G, Kline KG, Mohsin SB, Young JC, Otegui MS, Sussman MR. 2010. Molecular characterization of mutant Arabidopsis plants with reduced plasma membrane proton pump activity. Journal of Biological Chemistry 285: 17918–17929.

Haruta M, Gray WM, Sussman MR. 2015. Regulation of the plasma membrane proton pump (H^+^-ATPase) by phosphorylation. Current Opinion in Plant Biology 28: 68–75.

Hasson A, Plessis A, Blein T, Adroher B, Grigg S, Tsiantis M, Boudaoud A, Damerval C, Laufs P. 2011. Evolution and diverse roles of the CUP-SHAPED COTYLEDON genes in Arabidopsis leaf development. Plant Cell 23: 54–68.

Hattori T, Soga K, Wakabayashi K, Hoson T. 2022. An Arabidopsis *PTH2* gene is responsible for gravity resistance supporting plant growth under different gravity conditions. Life 12: 1603.

Hayashi Y, Nakamura S, Takemiya A, Takahashi Y, Shimazaki KI, Kinoshita T. 2010. Biochemical characterization of in vitro phosphorylation and dephosphorylation of the plasma membrane H^+^-ATPase. Plant and Cell Physiology 51: 1186–1196.

Hayashi Y, Takahashi K, Inoue SI, Kinoshita T. 2014. Abscisic acid suppresses hypocotyl elongation by dephosphorylating plasma membrane H^+^-ATPase in *Arabidopsis thaliana*. Plant and Cell Physiology 55: 845–853.

Hepworth J, Lenhard M. 2014. Regulation of plant lateral-organ growth by modulating cell number and size. Current Opinion in Plant Biology 17: 36–42.

Huang T, Irish VF. 2015. Temporal control of plant organ growth by TCP Transcription factors. Current Biology 25: 1765–1770.

Johansson F, Sommarin M, Larsson C. 1993. Fusicoccin activates the plasma membrane H^+^-ATPase by a mechanism involving the C-terminal inhibitory domain. Plant Cell 5: 321–327.

Koyama T, Furutani M, Tasaka M, Ohme-Takagi M. 2007. TCP transcription factors control the morphology of shoot lateral organs via negative regulation of the expression of boundary-specific genes in *Arabidopsis*. Plant Cell 19: 473–484.

Koyama T, Mitsuda N, Seki M, Shinozaki K, Ohme-Takagi M. 2010. TCP transcription factors regulate the activities of ASYMMETRIC LEAVES1 and miR164, as well as the auxin response, during differentiation of leaves in *Arabidopsis*. Plant Cell 22: 3574–3588.

Koyama T, Nii H, Mitsuda N, Ohta M, Kitajima S, Ohme-Takagi M, Sato F. 2013. A regulatory cascade involving class II ETHYLENE RESPONSE FACTOR transcriptional repressors operates in the progression of leaf senescence. Plant Physiology 162: 991– 1005.

Koyama T, Ohme-Takagi M, Sato F. 2011. Generation of serrated and wavy petals by inhibition of the activity of TCP transcription factors in *Arabidopsis thaliana*. Plant Signaling and Behavior 6: 697–699.

Koyama T, Sato F, Ohme-Takagi M. 2017. Roles of miR319 and TCP transcription factors in leaf development. Plant Physiology 175: 874–885.

Krahmer J, Fankhauser C. 2024. Environmental control of hypocotyl elongation. Annual Review of Plant Biology 75:489–519.

Kutscheras U. 1994. The current status of the acid-growth hypothesis. New Phytologist 126: 349–569.

Lan J, Qin G. 2020. The regulation of CIN-like TCP transcription factors. International Journal of Molecular Sciences 21: 4498.

Lan J, Wang N, Wang Y, Jiang Y, Yu H, Cao X, Qin G. 2023. Arabidopsis TCP4 transcription factor inhibits high temperature-induced homeotic conversion of ovules. Nature Communications 14: 5673

Li Y, Liu Z-B, Shi X, Hagen G, Guilfoyle TI. 1994. An auxin-inducible element in soybean SAUR promoters. Plant Physiology 106: 37–43.

Li L, Verstraeten I, Roosjen M, Takahashi K, Rodriguez L, Merrin J, Chen J, Shabala L, Smet W, Ren H, et al. 2021. Cell surface and intracellular auxin signalling for H+ fluxes in root growth. Nature 599: 273–277.

Li K, Yu R, Fan LM, Wei N, Chen H, Deng XW. 2016. DELLA-mediated PIF degradation contributes to coordination of light and gibberellin signalling in *Arabidopsis*. Nature Communications 7: 11867.

Lin W, Zhou X, Tang W, Takahashi K, Pan X, Dai J, Ren H, Zhu X, Pan S, Zheng H, et al. 2021. TMK-based cell-surface auxin signalling activates cell-wall acidification. Nature 599: 278–282.

Marowa P, Ding A, Kong Y. 2016. Expansins: roles in plant growth and potential applications in crop improvement. Plant Cell Reports 35: 949–965.

McQueen-Mason S, Durachko DM, Cosgrove DJ. 1992. Two endogenous proteins that induce cell wall extension in plants. Plant Cell 4: 1425–1433.

Miao R, Russinova E, Rodriguez PL. 2022. Tripartite hormonal regulation of plasma membrane H^+^-ATPase activity. Trends in Plant Science 27: 588–600.

Milani P, Gholamirad M, Traas J, Arnéodo A, Boudaoud A, Argoul F, Hamant O. 2011. In vivo analysis of local wall stiffness at the shoot apical meristem in Arabidopsis using atomic force microscopy. Plant Journal 67: 1116–1123.

Minami A, Takahashi K, Inoue Sichiro, Tada Y, Kinoshita T. 2019. Brassinosteroid induces phosphorylation of the plasma membrane H^+^-ATPase during hypocotyl elongation in *Arabidopsis thaliana*. Plant and Cell Physiology 60: 935–944.

Mitsuda N, Hiratsu K, Todaka D, Nakashima K, Yamaguchi-Shinozaki K, Ohme-Takagi M. 2006. Efficient production of male and female sterile plants by expression of a chimeric repressor in *Arabidopsis* and rice. Plant Biotechnology Journal 4: 325–332.

Nagpal P, Reeves PH, Wong JH, Armengot L, Chae K, Rieveschl NB, Trinidad B, Davidsdottir V, Jain P, Gray WM, et al. 2022. SAUR63 stimulates cell growth at the plasma membrane. PLoS Genetics 18: e1010375.

Nath U, Crawford BCW, Carpenter R, Coen E. 2003. Genetic control of surface curvature. Science 299: 1404–1407.

Ni M, Tepperman JM, Quail PH. 1998. PIF3, a phytochrome-interacting factor necessary for normal photoinduced signal transduction, is a novel basic helix-loop-helix protein. Cell 95: 657–667.

Nicolas M, Cubas P. 2016. TCP factors: New kids on the signaling block. Current Opinion in Plant Biology 33: 33–41.

Okumura M, Inoue Sichiro, Kuwata K, Kinoshita T. 2016. Photosynthesis activates plasma membrane H^+^-ATPase via sugar accumulation. Plant Physiology 171: 580–589.

Palatnik JF, Allen E, Wu X, Schommer C, Schwab R, Carrington JC, Weigel D. 2003. Control of leaf morphogenesis by microRNAs. Nature 425: 257–263.

Peaucelle A, Braybrook SA, Le Guillou L, Bron E, Kuhlemeier C, Höfte H. 2011. Pectin-induced changes in cell wall mechanics underlie organ initiation in *Arabidopsis*. Current Biology 21: 1720–1726.

Peaucelle A, Wightman R, Höfte H. 2015. The control of growth symmetry breaking in the *Arabidopsis* hypocotyl. Current Biology 25: 1746–1752.

Pien SP, Wyrzykowska J, Mcqueen-Mason S, Smart C, Fleming A. 2001. Local expression of expansin induces the entire process of leaf development and modifies leaf shape. Proceedings of National Academy of Science of USA 98: 11812–11817.

Rath M, Challa KR, Sarvepalli K, Nath U. 2022. CINCINNATA-Like TCP transcription factors in cell growth – An expanding portfolio. Frontiers in Plant Science 13: 825341.

Ren H, Gray WM. 2015. SAUR proteins as effectors of hormonal and environmental signals in plant growth. Molecular Plant 8: 1153–1164.

Rose JKC, Braam J, Fry SC, Nishitani K. 2002. The XTH family of enzymes involved in xyloglucan endotransglucosylation and endohydrolysis: Current perspectives and a new unifying nomenclature. Plant and Cell Physiology 43: 1421–1435.

Rubio-Somoza I, Zhou CM, Confraria A, Martinho C, Von Born P, Baena-Gonzalez E, Wang JW, Weigel D. 2014. Temporal control of leaf complexity by miRNA-regulated licensing of protein complexes. Current Biology 24: 2714–2719.

Rui Y, Dinneny JR. 2020. A wall with integrity: surveillance and maintenance of the plant cell wall under stress. New Phytologist 225: 1428–1439.

Rayle D, Cleland R. 1992. The acid growth theory of auxin-induced cell elongation is alive and well. Plant Physiology 99: 1271–1274.

Sarvepalli K, Das Gupta M, Challa KR, Nath U. 2019. Molecular cartography of leaf development — role of transcription factors. Current Opinion in Plant Biology 47: 22– 31.

Schindelin J, Arganda-Carreras I, Frise E, Kaynig V, Longair M, Pietzsch T, Preibisch S, Rueden C, Saalfeld S, Schmid B, et al. 2012. Fiji: An open-source platform for biological-image analysis. Nature Methods 9: 676–682.

Schneider M, Van Bel M, Inzé D, Baekelandt A. 2024. Leaf growth – complex regulation of a seemingly simple process. Plant Journal 117: 1018–1051.

Schommer C, Debernardi JM, Bresso EG, Rodriguez RE, Palatnik JF. 2014. Repression of cell proliferation by miR319-regulated TCP4. Molecular Plant 7: 1533– 1544.

Shani E, Salehin M, Zhang Y, Sanchez SE, Doherty C, Wang R, Mangado CC, Song L, Tal I, Pisanty O, et al. 2017. Plant stress tolerance requires auxin-sensitive Aux/IAA transcriptional repressors. Current Biology 27: 437–444.

Shankar N, Sunkara P, Nath U. 2023. A double-negative feedback loop between miR319c and JAW-TCPs establishes growth pattern in incipient leaf primordia in *Arabidopsis thaliana*. PloS Genetics 19: e1010978.

Shigeyama T, Watanabe A, Tokuchi K, Toh S, Sakurai N, Shibuya N, Kawakami N. 2016. α-Xylosidase plays essential roles in xyloglucan remodelling, maintenance of cell wall integrity, and seed germination in *Arabidopsis thaliana*. Journal of Experimental Botany 67: 5615–5629.

Spartz AK, Ren H, Park MY, Grandt KN, Lee SH, Murphy AS, Sussman MR, Overvoorde PJ, Gray WM. 2014. SAUR inhibition of PP2C-D phosphatases activates plasma membrane H^+^-ATPases to promote cell expansion in *Arabidopsis*. Plant Cell 26: 2129–2142.

Storey JD. 2002. A direct approach to false discovery rates. *Journal of Royal Statistical Society*, Series B 64:479–498.

Takahashi K, Hayashi KI, Kinoshita T. 2012. Auxin activates the plasma membrane H^+^-ATPase by phosphorylation during hypocotyl elongation in *Arabidopsis*. Plant Physiology 159: 632–641.

Tao Q, Guo D, Wei B, Zhang F, Pang C, Jiang H, Zhang J, Wei T, Gu H, Qu LJ, et al. 2013. The TIE1 transcriptional repressor links TCP transcription factors with TOPLESS/TOPLESS-RELATED corepressors and modulates leaf development in *Arabidopsis*. Plant Cell 25: 421–437.

Tian, JQ, Reed W. 1999. Control of auxin-regulated root development by the *Arabidopsis thaliana* SHY2/IAA3 gene. Development 126: 711–712.

Trigg SA, Garza RM, MacWilliams A, Nery JR, Bartlett A, Castanon R, Goubil A, Feeney J, O’Malley R, Huang SSC, et al. 2017. CrY2H-seq: A massively multiplexed assay for deep-coverage interactome mapping. Nature Methods 14: 819–825.

Trinh DC, Alonso-Serra J, Asaoka M, Colin L, Cortes M, Malivert A, Takatani S, Zhao F, Traas J, Trehin C, et al. 2021. How mechanical forces shape plant organs. Current Biology 31: R143–R159.

Tsukaya H. 2021. The leaf meristem enigma: The relationship between the plate meristem and the marginal meristem. Plant Cell 33: 3194–3206.

Tsukaya H, Tsuge T, Uchimiya H. 1994. The cotyledon: a superior system for studies of leaf development. Planta 195: 309–312.

Vanhaeren H, Gonzalez N, Coppens F, De Milde L, Van Daele T, Vermeersch M, Eloy NB, Storme V, Inzé D. 2014. Combining growth-promoting genes leads to positive epistasis in *Arabidopsis thaliana*. eLife 3: e02252.

Wei B, Zhang J, Pang C, Yu H, Guo D, Jiang H, Ding M, Chen Z, Tao Q, Gu H, et al. 2015. The molecular mechanism of SPOROCYTELESS/NOZZLE in controlling Arabidopsis ovule development. Cell Research 25: 121–134.

Wu Q, Li Y, Lyu M, Luo Y, Shi H, Zhong S. 2020. Touch-induced seedling morphological changes are determined by ethylene-regulated pectin degradation. Science Advances 6: eabc9294.

Xu N, Hagen G, Guilfoyle T. 1997. Multiple auxin response modules in the soybean SAUR 15A promoter. Plant Science 126:193–201.

Zheng X, Lan J, Yu H, Zhang J, Zhang Y, Qin Y, Su XD, Qin G. 2022. Arabidopsis transcription factor TCP4 represses chlorophyll biosynthesis to prevent petal greening. Plant Communications 3: 100309.

Zhou Y, Xun Q, Zhang D, Lv M, Ou Y, Li J. 2019. TCP Transcription factors associate with PHYTOCHROME INTERACTING FACTOR 4 and CRYPTOCHROME 1 to regulate thermomorphogenesis in *Arabidopsis thaliana*. iScience 15: 600–610.

Zuch DT, Doyle SM, Majda M, Smith RS, Robert S, Torii KU. 2022. Cell biology of the leaf epidermis: Fate specification, morphogenesis, and coordination. Plant Cell 34: 209–227.

Zuo J, Niu Q-W, Chua N-H. 2000. An estrogen receptor-based transactivator XVE mediates highly inducible gene expression in transgenic plants. Plant Journal. 24: 265– 273.

